# FoxA1/2-dependent epigenomic reprogramming drives lineage switching in lung adenocarcinoma

**DOI:** 10.1101/2023.10.30.564775

**Authors:** Katherine Gillis, Walter A. Orellana, Emily Wilson, Timothy J. Parnell, Gabriela Fort, Headtlove Essel Dadzie, Xiaoyang Zhang, Eric L. Snyder

**Affiliations:** Huntsman Cancer Institute, University of Utah, Salt Lake City, UT, USA; Department of Oncological Sciences, University of Utah, Salt Lake City, UT, USA; Department of Pathology, University of Utah, Salt Lake City, UT, USA

**Keywords:** lung adenocarcinoma, FoxA1/2, TET3, cancer lineage switching/ plasticity, epigenetic reprogramming

## Abstract

The ability of cancer cells to alter their identity is essential for tumor survival and progression. Loss of the pulmonary lineage specifier NKX2-1 within KRAS-driven lung adenocarcinoma (LUAD) enhances tumor progression and results in a pulmonary-to-gastric lineage switch that is dependent upon the activity of pioneer factors FoxA1 and FoxA2; however, the underlying mechanism remains largely unknown. Here, we show that FoxA1/2 reprogram the epigenetic landscape of NKX2-1-negative LUAD to facilitate a gastric identity. After *Nkx2-1* deletion, FoxA1/2 mediate demethylation of gastric-defining genes through recruitment of TET3, an enzyme that induces DNA demethylation. H3K27ac ChIP-seq and HiChIP show that FoxA1/2 also control the activity of regulatory elements and their 3D interactions at gastric loci. Furthermore, oncogenic KRAS is required for the FoxA1/2-dependent epigenetic reprogramming. This work demonstrates the role of FoxA1/2 in rewiring the methylation and histone landscape and cis-regulatory dynamics of NKX2-1-negative LUAD to drive cancer cell lineage switching.

## INTRODUCTION

Cellular identity is a fundamental aspect of normal tissue development and homeostasis, dictating cell function and fate within multicellular organisms. The ability of cells to alter their identity, a process known as lineage switching/plasticity or transdifferentiation, is essential during embryonic development and tissue repair^1^. However, in the context of cancer, this highly regulated process can be exploited by tumor cells, allowing them to adopt alternative cell fates that provide a selective advantage^2^. There are several contexts under which cancer cell lineage switching occurs including: (i) natural selection as a response to microenvironmental constraints or the acquisition of genetic alterations^2,3^; (ii) acute drug response resulting in the formation of drug-tolerant persister populations^4,5^; and (iii) secondary drug response as a mechanism of acquired resistance to therapy^6,7^. Thus, there is a clear need to better understand the molecular mechanisms that regulate lineage switching, with the ultimate goal of identifying potential therapeutic strategies to either counteract these changes or target novel identity-specific vulnerabilities.

Cell identity can be enforced through multiple epigenetic mechanisms. First, DNA methylation is a stable epigenetic mark required for the establishment and maintenance of cell identity. Lineage-specific methylation patterns are necessary to preserve fidelity of transcriptional programs and reinforce cell type-specific characteristics^8–10^. Aberrant DNA methylation, characterized by global hypomethylation and site-specific hypermethylation, is a hallmark of cancer cells^11,12^. Such epigenetic alterations can lead to dysregulation of gene expression (e.g., DNA hypermethylation-mediated silencing of tumor suppressor genes) and facilitate tumor initiation and progression. In addition to DNA methylation, histone post-translational modifications (PTMs) and 3D chromatin structure have also been linked to the regulation of cell identity^13,14^. During embryonic stem cell differentiation, cis-regulatory interactions and histone PTMs are globally remodeled to facilitate expression of lineage-specific genes^15,16^. Epigenetic dysregulation via altered enhancer-promoter interactions or impaired histone deposition has been shown to directly alter cancer cell identity and drive tumor progression^17^. Although the association between a cell’s epigenome and its identity is well established, the molecular mechanisms responsible for dictating chromatin dynamics is not as well understood.

Lung adenocarcinoma (LUAD), the most frequently diagnosed subtype of lung cancer, exhibits substantial heterogeneity in its cellular identity. Invasive mucinous adenocarcinoma (IMA) is a subtype of LUAD (∼5-10% of cases) that undergoes pulmonary to gastric lineage switching during its natural progression. Previous work using genetically engineered mouse models (GEMMs) has shown that loss of the pulmonary lineage specifying transcription factor (TF), NKX2-1/TTF1, causes gastric transdifferentiation in LUAD, generating murine tumors that recapitulate the morphology and gene expression profile of human IMA^18^. Although ∼75% of human LUAD express NKX2-1, IMAs typically lose NKX2-1 expression due to point mutations or epigenetic silencing^19^. Moreover, ∼75% of IMAs are driven by oncogenic KRAS (vs. 20-30% of LUAD overall)^20^. As a result, targeted therapy for IMA has lagged behind other LUAD subtypes driven by oncogenic kinases (such as *EGFR*) that can be inhibited by small molecules^20^.

We have previously shown that the pulmonary to gastric lineage switch caused by NKX2-1 loss in LUAD is mediated by differential chromatin binding of the forkhead box TFs, FoxA1 and FoxA2. The FoxA proteins are expressed in a variety of endodermal tissues and have important roles in development, differentiation, tumorigenesis, and metabolism^21^. In addition to their TF activity, FoxA1/2 can modulate the epigenetic landscape by multiple mechanisms. FoxA1/2 are known to open chromatin structure via pioneer factor activity^21^, alter histone post-translational modifications (PTMs) through direct recruitment of histone-modifying enzymes^22,23^, and facilitate DNA demethylation via interactions with the ten-eleven translocation (TET) dioxygenases^24^. In LUAD, FoxA1/2 colocalize with NKX2-1 at adjacent sites within the genome of both human^25^ and murine tumors^18^. These sites include regulatory elements for genes involved in pulmonary differentiation. Upon *Nkx2-1* deletion, FoxA1/2 lose binding at approximately half of these shared sites and relocate throughout the genome to de novo binding sites at regulatory elements of gastric genes including *Hnf4a*^18^. Many of these de novo binding sites are known targets of FoxA1/2 within the gastrointestinal tract^26^. *Nkx2-1* deletion also induces histone PTMs associated with gene activation at de novo FoxA1/2 binding sites, including an increase in histone 3 lysine 27 acetylation (H3K27ac). However, it is unknown whether FoxA1/2 directly mediate these chromatin modifications, or whether chromatin modifications occur independently of FoxA1/2 binding.

Here we investigate the epigenetic basis of gastric lineage switching in LUAD, shedding light on the role of specific TFs and epigenetic modifications in controlling cell identity during cancer progression. We employ sequential in vivo recombination systems to discern the precise role of FoxA1/2 in modifying the methylation and histone landscape of LUAD, as well as recruitment of methylation modifying enzymes. Using this system, we show that FoxA1/2 are responsible for TET3 recruitment and demethylation at lineage-specific sites in NKX2-1-negative LUAD. Additionally, we find that FoxA1/2 facilitate H3K27ac deposition and enhancer-promoter (E-P) interactions at genes defining gastric lineage. Finally, we show that oncogenic KRAS fine tunes the epigenetic state of LUAD, modifying the precise identity adopted by cells after NKX2-1 loss.

## RESULTS

### NKX2-1 loss leads to widespread DNA methylation changes in lung adenocarcinoma

DNA methylation is a stable epigenetic mark that governs cellular identity, with unique cell type-specific unmethylated regions delineating cell origin and type^8,27^. In numerous contexts, manipulation of the enzymes responsible for controlling DNA methylation has been shown to block cell differentiation^9,28,29^. Given this association, we asked whether the pulmonary-to-gastric lineage switch induced by NKX2-1 loss is accompanied by alterations in DNA methylation. To answer this question, we developed a sequential recombination GEMM that enables isolation of tumor cells and nuclei after *Nkx2-1* deletion in established KRAS-driven LUAD. We generated *Kras^FSF-G12D/+^*; *Rosa26^FSF-CreERT2/Sun1^; Nkx2-1^F/F^* (KN) mice as well as control mice harboring a single conditional allele of *Nkx2-1* (K). Tumors were initiated via intratracheal delivery of adenovirus expressing SPC-driven FlpO recombinase^30^, which activates oncogenic Kras^G12D^ and tamoxifen-inducible Cre^ERT2^ in the distal lung epithelium. Tamoxifen administration led to Cre^ERT2^-mediated deletion of *Nkx2-1* as well as expression of a GFP-tagged nuclear membrane protein, *Sun1* (Figure S1A). Both nuclei and cells were extracted from control and experimental tumors and FACS sorted for GFP-positivity to obtain a tumor-specific population (Figure S1B-D). DNA and RNA were isolated from sorted nuclei and cells respectively to evaluate methylation and transcriptional changes after *Nkx2-1* deletion.

Bulk RNA-seq on sorted tumor cells from K and KN tumors (n=3 replicates per genotype) identified 5,463 differentially expressed genes (DEGs; log_2_FC > 1; padj < 0.05, Table S1). Consistent with our previous findings^18^, loss of NKX2-1 within established LUAD tumors resulted in a pulmonary to gastric lineage switch (Figure S1E). Specifically, tumor cells shed their pulmonary identity and adopted a state similar to gastric pit cells, which line the lumen of the stomach. This is consistent with our previous observation that high levels of RAF/ERK activity promotes a gastric pit program in NKX2-1-negative LUAD^4^. Following validation of the gastric lineage switch, we then sought to profile the methylation changes that accompany NKX2-1 loss.

Whole-genome enzymatic methyl-seq (EM-seq) on sorted nuclei from K and KN tumors (n=3 replicates per genotype) demonstrated that NKX2-1 loss leads to extensive changes in DNA methylation throughout the LUAD genome. We identified 23,942 differentially methylated regions (DMRs) between K and KN tumors using the bsseq^31^ pipeline with a stringency cutoff of 15% difference in fraction methylation (Figure 1A and S2A; Table S2). Of these regions, 10,676 sites showed a significant gain in methylation with *Nkx2-1* deletion (“hyperDMRs”) while 13,266 showed a significant reduction in methylation (“hypoDMRs”). The NKX2-1 binding motif was the most highly enriched in hyperDMRs, suggesting that NKX2-1 maintains a hypomethylated state at many of its binding sites in K tumors (Figure 1B). Additionally, hyperDMRs exhibited a strong enrichment for motifs bound by TFs that promote alveolar type II (AT2) and type I (AT1) identity (CEBP and TEAD families respectively^32^). In contrast, hypoDMRs were enriched for motifs bound by TFs that can control gastrointestinal (GI) identity, including HNF4, GATA, and KLF. FoxA1/2-bound motifs were enriched in both hyper- and hypoDMRs, reflecting their ability to promote pulmonary or gastric identity in LUAD^18,30^. Of note, the NKX motif was not enriched in hypoDMRs. This is consistent with our prior report that NKX2-1 inhibits gastric marker gene transcription predominantly by controlling FoxA1/2 localization rather than direct binding^18^.

**Figure 1:**
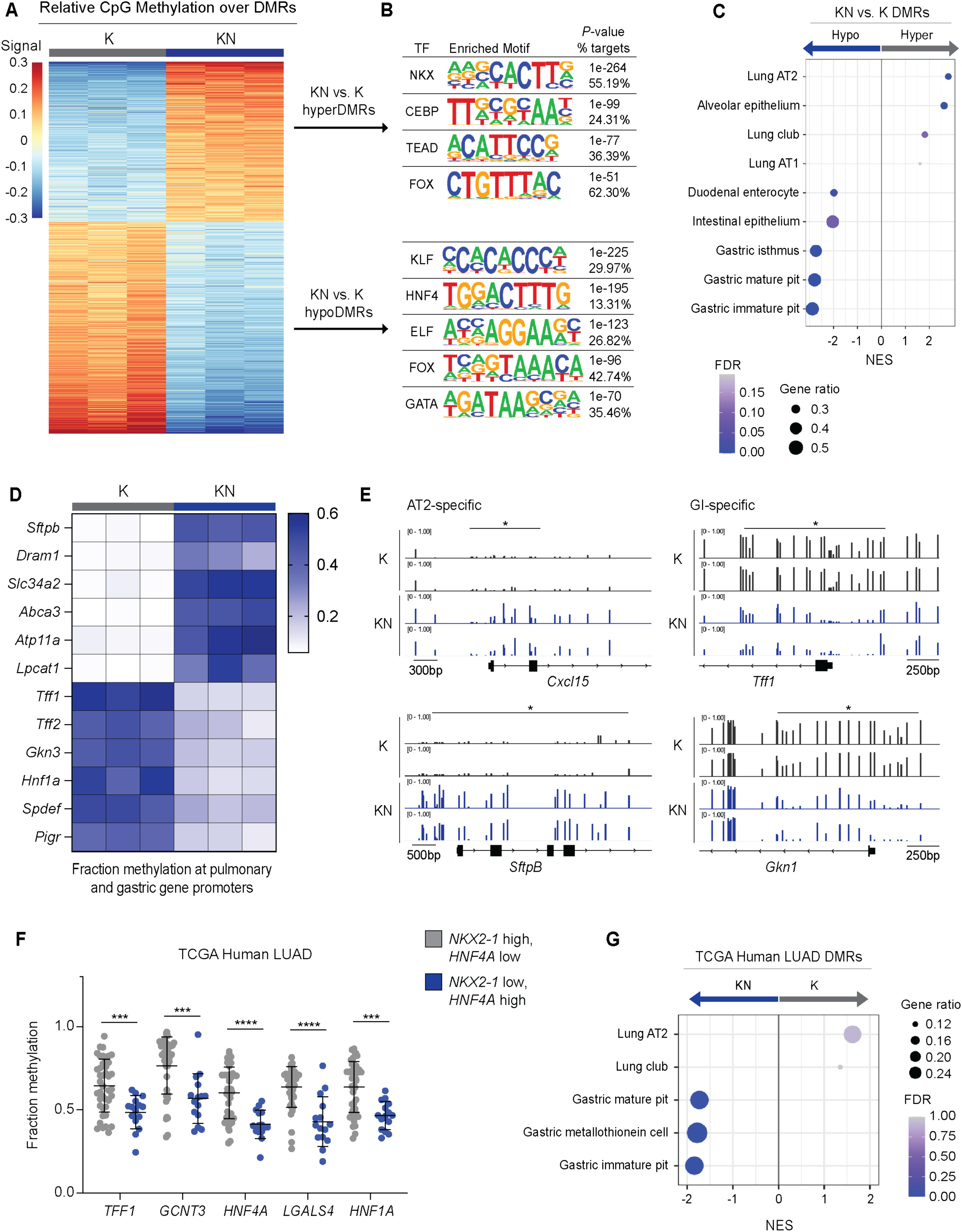
NKX2-1 loss leads to widespread DNA methylation changes in lung adenocarcinoma. A) Heatmap of DMRs identified in K and KN tumors showing relative difference in mean methylation at CpGs. DNA methylation scores calculated by subtracting the row mean from the average methylation level of each DMR per sample. Samples collected 14-weeks post tumor initiation. B) TF motifs enriched in hyper- and hypoDMRs identified between K and KN samples. C) GSEA for cell type signatures on DMGs identified between K and KN samples. D) Heatmap showing the average fraction methylation values at DMRs localized to pulmonary and gastric gene promoters in K and KN samples. E) Methylation tracks of AT2 (left) or GI (right) genes in K and KN tumors. Lines with asterisk indicate significant DMRs. F) Fraction methylation values at GI-specific promoters in KRAS-driven human LUAD categorized by *NKX2-1* and *HNF4a* expression (TCGA Pan Cancer Atlas). G) GSEA for cell type signatures on DMGs identified from K and KN human LUAD (TCGA Pan Cancer Atlas).

To identify biological processes and functions regulated by these global methylation changes, we annotated DMRs to their adjacent genes, which identified 3,362 hyper- and 4,122 hypomethylated genes in KN vs. K tumors. Gene Set Enrichment Analysis (GSEA) of these differentially methylated genes (DMGs) revealed an enrichment for pulmonary and gastric cell type signatures in hyper- and hypomethylated genes respectively (Figure 1C). Signatures of gastric pit cells were the most significantly enriched among hypomethylated genes. Enrichr analysis with the Genotype-Tissue expression (GTEx) database substantiated these lineage-specific methylation changes, demonstrating an enrichment for lung and gastric tissue types (Table S3). Fraction methylation profiles of DMRs located at pulmonary and gastric gene promoters highlight their contrasting methylation changes following NKX2-1 loss (Figure 1D). Numerous AT2 marker genes, including *Sftpb*, *Cxcl15 and Slc34a2* exhibited a significant increase in methylation following *Nkx2-1* deletion (Figure 1E). The canonical AT2 marker *Sftpc* exhibited a noticeable increase in methylation, although it did not meet the significance threshold of 15%. Of note, a subset of AT2 markers, including *Napsa* and *Sftpd,* remained hypomethylated at their promoters even in a gastric state (Figure S2B). Conversely, markers of gastric pit cells (e.g., *Tff1* and *Gkn1*), as well as pan gastric markers (e.g., *Hnf4a* and *Lgals4*) demonstrated a significant decrease in their fraction methylation. However, genes expressed in the distal GI tract (e.g., *Cdx2* and *Muc2*), which are not induced upon *Nkx2-1* deletion, did not undergo significant methylation changes.

Intersection of methylation and transcriptional datasets revealed a substantial overlap: 34.0% of upregulated genes after *Nkx2-1* deletion showed a significant decrease in methylation (versus 11.8% of downregulated genes). Conversely, 24.4% of downregulated genes had a corresponding increase in methylation (versus 12.3% of upregulated genes). Importantly, many AT2 and gastric-specific marker genes demonstrated concomitant methylation and transcriptional changes, suggesting that DNA methylation may be one epigenetic mechanism regulating expression of lineage-defining genes in LUAD.

We next sought to understand the extent to which DMRs induced by *Nkx2-1* deletion in GEMMs correlate with DNA methylation patterns in normal human cells in vivo. First, we assessed the magnitude of DMRs between human lung and GI epithelial cells profiled in the human methylome atlas^8^. Using the same analysis pipeline and thresholds, we found that fully differentiated alveolar and gastric epithelial cells contain 73,977 DMRs (Table S2). Therefore, loss of single TF within KRAS-driven LUAD leads to approximately one third as many DMRs as two developmentally distinct cell fate trajectories. Next, we intersected the top 1000 differentially unmethylated regions in human alveolar and gastric epithelial cell types with DMRs identified after *Nkx2-1* deletion. We found that hypoDMRs were significantly more associated with gastric unmethylated regions while hyperDMRs correlated more with alveolar unmethylated regions (Fisher’s exact test; *p*-val<0.05). Finally, we evaluated TF motifs associated with cell-type-specific unmethylated regions defined by the human methylome atlas^8^. Intriguingly, NKX2-1 and FOX motifs were the most highly enriched in normal lung alveolar cells, corresponding with our motif analysis for hyperDMRs. This suggests that in LUAD, similar to human alveolar cells, NKX2-1 and FoxA1/2 promote a pulmonary identity by mediating demethylation of AT2-specific sites. Conversely, motifs enriched in hypoDMRs, including HNF4, GATA, and KLF families, were significantly associated with unmethylated regions in human GI tissues including gastric, GI, and small intestine cell types. These data support a model in which TFs regulate LUAD cell identity via methylation reprogramming that reflects their normal epigenetic function.

Finally, we asked whether similar lineage-specific DNA methylation patterns occurred in human LUAD. To answer this, *KRAS*-mutant LUAD samples from the TCGA PanCancer Atlas (n= 154/562) were categorized into two groups: *NKX2-1*-high, *HNF4A*-low (i.e., K tumors; n=41, Table S1) and *NKX2-1*-low, *HNF4A*-high (i.e., KN tumors; n=15, Table S1). GSEA on DEGs (n=1,806; log_2_FC>1; padj<0.05, Table S2) between the two groups demonstrated significant enrichment for alveolar signatures in human K tumors, whereas human KN samples were most highly enriched for gastric pit cell signatures (Figure S2D). Evaluation of DMRs (n=1,780; padj<0.05; qval<0.25) revealed that many pan-gastric and pit cell marker genes had significantly lower DNA methylation levels in human KN samples compared to K tumors (Figure 1F). In fact, the top enriched cell type signature for KN hypomethylated genes was the gastric immature pit cell (Figure 1G). Pulmonary AT2 markers were heterogenous, as some genes had higher DNA methylation in K tumors (e.g., *SFTPB* and *NAPSA*) whereas others showed no significant differences (e.g., *SFTPC*). Altogether, these data suggest that stochastic NKX2-1 loss during human LUAD progression and deletion of *Nkx2-1* in LUAD GEMMs both cause similar, highly conserved changes in DNA methylation patterns.

### FoxA1/2 are required to demethylate gastric gene regulatory elements to facilitate the pulmonary-to-gastric lineage switch in LUAD

Several recent studies have found that FoxA1/2 control cellular identity by facilitating DNA demethylation at lineage-defining genes both *in vitro* and *in vivo*^24,29^. These demethylation events are dependent on the activity of TET dioxygenases, the enzymes that mediate DNA demethylation through progressive oxidative reactions at methyl-cytosines^33^. FoxA1/2-mediated demethylation is necessary for enhancer activation and gene expression of lineage-specific targets. We therefore investigated whether FoxA1/2 facilitate the profound DNA methylation changes that occur after NKX2-1 loss in LUAD. We generated a GEMM that concomitantly deleted *Foxa1/2* and *Nkx2-1* in established KRAS-driven neoplasia (KNF1F2) and performed bulk RNA-seq on sorted KNF1F2 tumor cells (Figure S3A). This identified 4,077 DEGs resulting from *Foxa1/2* deletion in KN tumors (log_2_FC > 1; padj < 0.05, Table S1). Consistent with our previous findings^30^, *Foxa1/2* deletion *in vivo* completely prevented gastric transdifferentiation (Figure S3B-C). Instead, KNF1F2 tumors adopt either a squamous state or a state resembling the squamocolumnar junction (SCJ) of the stomach^30^. In addition to cell identity changes, FoxA1/2 loss also resulted in downregulation of processes involved in cholesterol, bile acid, and fatty acid metabolism (Figure S3D). Of note, FoxA1/2 loss within established KN tumors did not significantly alter tumor burden as we were able to sort similar numbers of cells and nuclei from both genotypes (approximately 80 – 100 million per KN or KNF1F2 lung; ∼10 fold higher than K controls).

Given the necessity of FoxA1/2 for establishment of a gastric program, we next investigated their role in DNA methylation by performing whole genome EM-seq on sorted nuclei from KNF1F2 tumors (Figure 2A and S3E, Table S2). To identify FoxA1/2 dependent methylation changes, DMRs were merged from distinct pairwise comparisons between the tumor types (as described in “DNA Methylation Sequencing” section of methods). We first evaluated regions demethylated after *Nkx2-1* deletion. Of the 10,355 hypoDMRs, the vast majority (80%) were significantly higher in KNF1F2 than KN, showing that their demethylation was FoxA1/2 dependent (Figure 2A, purple). Motif analysis of these FoxA1/2 dependent hypoDMRs showed an enrichment for TFs that promote GI identity including HNF4, FOX, and SPDEF families (Figure 2B). Annotation and gene ontology (GO) analysis revealed that genes demethylated in a FoxA1/2 dependent manner were most significantly enriched for gastric pit and isthmus cell signatures (Figure 2C). Fraction methylation profiles of DMRs located at multiple pan-gastric and pit cell marker gene promoters demonstrate a striking dependency on FoxA1/2 for demethylation (Figure 2D; representative methylation tracks at gastric pit cell markers, *Tff1* and *Gkn1* depicted in Figure 2E and S3F). Altogether, these data show that FoxA1/2 are required for demethylation of lineage-defining sites during gastric transdifferentiation in LUAD.

**Figure 2:**
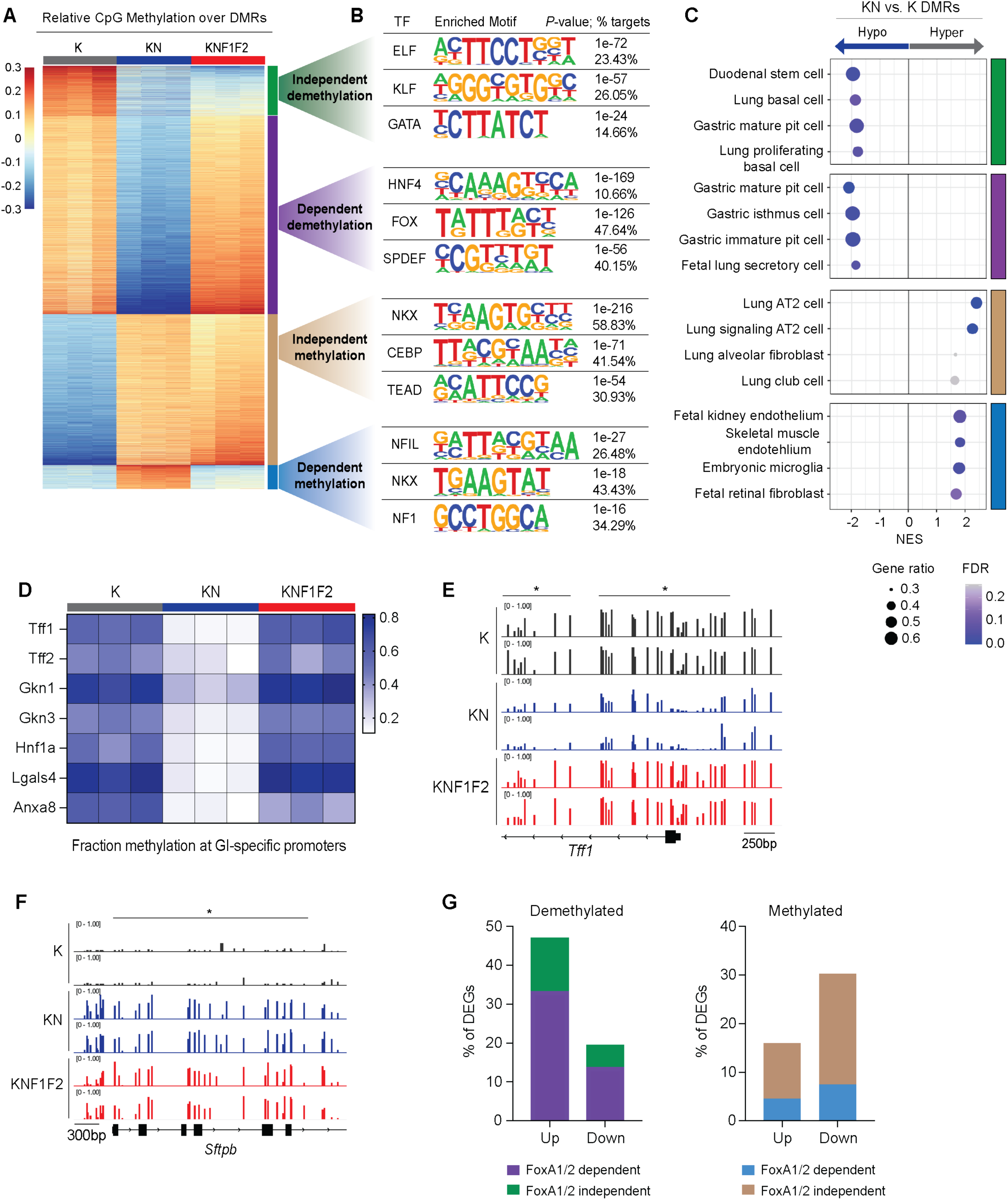
FoxA1/2 are required to demethylate gastric gene regulatory elements to facilitate the pulmonary-to-gastric lineage switch in LUAD. A) Heatmap showing relative difference in mean methylation of CpGs at DMRs identified in K, KN, and KNF1F2 tumors. DMRs categorized by FoxA1/2-dependency and methylation status (e.g., methylated vs. demethylated). Samples collected 14-weeks post tumor initiation. B) TF motifs enriched in FoxA1/2 dependent vs. independent methylated and demethylated regions following *Nkx2-1* deletion. C) GSEA for cell type signatures on DMGs identified from each of the four categories defined in Figure 2B. D) Heatmap showing average fraction methylation values at DMRs localized to gastric gene promoters in K, KN, and KNF1F2 tumors. E) Methylation tracks at gastric pit cell gene, *Tff1*, in K, KN, and KNF1F2 tumors. Lines with asterisk indicate significant DMRs. F) Methylation tracks at AT2 cell gene, *Sftpb*, in K, KN, and KNF1F2 tumors. Lines with asterisk indicate significant DMRs. G) Intersection of KN vs. K DEGs with DMRs defined in Figure 2A. Regions demethylated in KN tumors, regardless of FoxA1/2 dependency, are primarily associated with upregulated genes. Conversely, methylated regions are associated with downregulated genes.

In contrast, regions demethylated in a FoxA1/2-independent manner following *Nkx2-1* deletion (Figure 2A, green) showed no enrichment for FOX and HNF4 motifs, but a significant association with other TF’s involved in epithelial and GI differentiation (e.g., KLF and GATA; Figure 2B). GSEA of FoxA1/2 independent hypoDMRs revealed an enrichment for both basal and GI cell type signatures. Enrichr analysis using the GTEx database showed that these regions correlate most significantly with human esophageal tissue (Table S3), which is consistent with the presence of squamous cell carcinomas (SCCs) in KNF1F2 mice. Given that gastric differentiation is dependent on FoxA1/2 at the transcriptomic and morphologic level, we were surprised to find pit cell and duodenal signatures enriched for genes demethylated in a FoxA1/2-independent manner. We therefore asked whether demethylation of these GI genes leads to increased expression, even in KNF1F2 samples. First, we examined the methylation status of the 27 genes driving the gastric pit cell signature and found that all but one contained both FoxA1/2 dependent and independent demethylation sites. FoxA1/2-independent sites also tended to be located further away from the gene itself and were comparatively less abundant. Intersection with bulk RNA-seq showed that these methylation changes correlated strikingly well with gene expression. Genes comprising the gastric pit cell signature demonstrated the highest expression in KN tumors; however, these genes were induced to a lesser extent in KNF1F2 samples compared to K controls (Figure S3G). Of note, some of these genes (e.g., *Klf3/4*) can also play a role in squamous differentiation, suggesting that they may not be as specific to pit cells as some canonical markers, such as *Gkn1* and *Tff1*^34,35^. Thus, there are FoxA1/2-independent mechanisms that partially demethylate and transcriptionally activate a subset of genes in the pit cell signature upon *Nkx2-1* deletion. However, FoxA1/2 are required for full demethylation and activation of a gastric pit cell program in NKX2-1-negative LUAD, particularly the most specific marker genes such as *Gkn1* and *Tff1*.

We then asked whether hyperDMRs are also FoxA1/2 dependent. We found that 84.5% of the 7,212 hyperDMRs did not require FoxA1/2. Motif analysis of FoxA1/2 independent hyperDMRs revealed an enrichment for TFs that control pulmonary identity, including NKX, CEBP, and TEAD families (Figure 2B; beige). These findings correspond with GO analysis showing that genes methylated in a FoxA1/2 independent manner were significantly enriched for AT2 cell signatures (Figure 2C). Evaluation of individual genes revealed that most AT2-specific sites gained methylation after NKX2-1 loss irrespective of FoxA1/2 activity including canonical targets *Sftpb*, *Cxcl15*, and *Dram1* (Figure 2F and data not shown).

FoxA1/2-dependent hyperDMRs comprised the smallest number of DMRs (n=1,113 sites). Although these regions were enriched for the NKX motif, the FOX motif was not enriched, suggesting that their FoxA1/2-dependency is more indirect than FoxA1/2-dependent hypoDMRs (Figure 2B; blue; see also Figure 3 for analysis of FoxA1 genomic localization). GO analysis on FoxA1/2 dependent hyperDMRs did not show a consistent enrichment for a specific cell type, perhaps due to the small number of sites.

**Figure 3:**
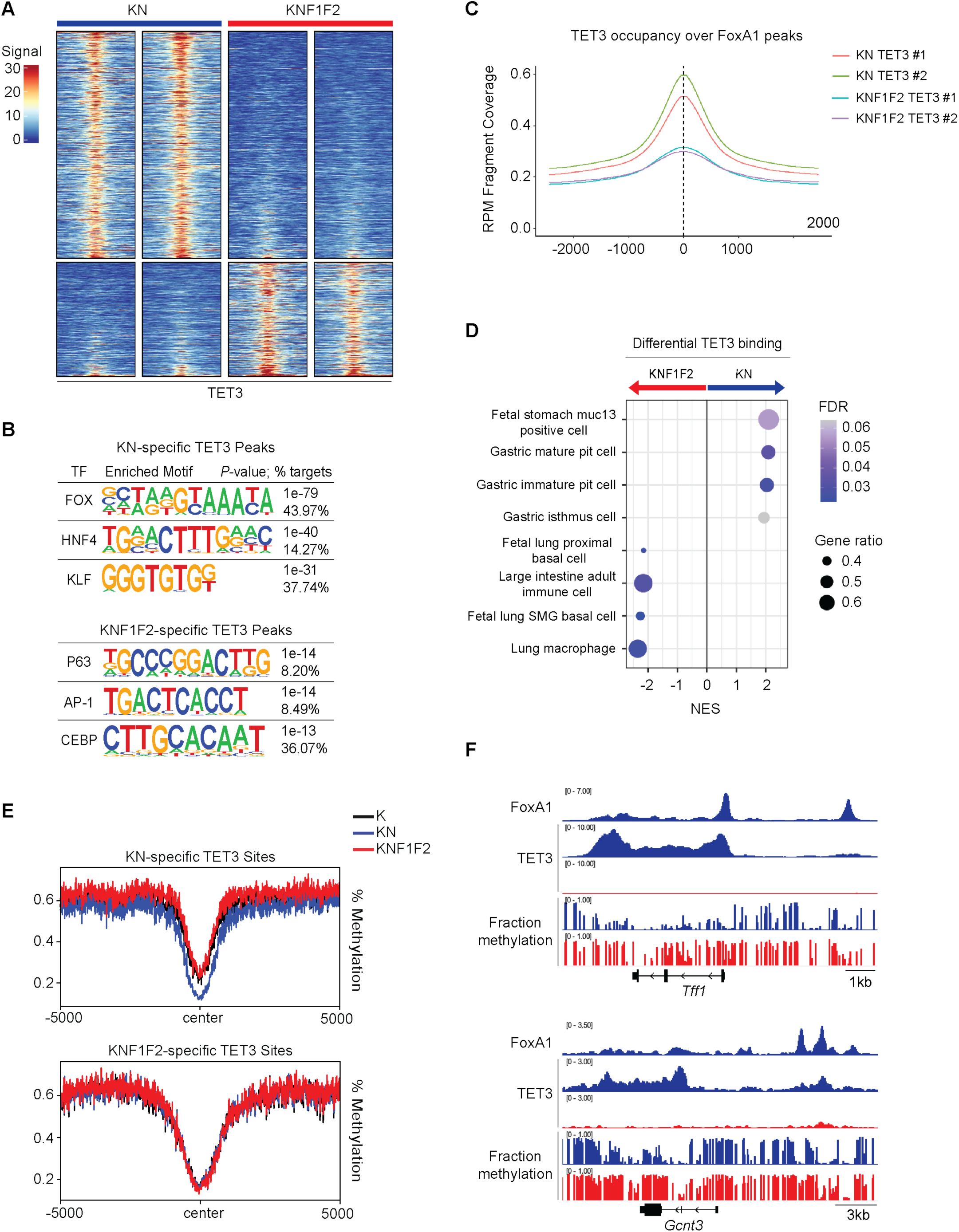
FoxA1/2 recruit TET3 to lineage-defining gastric sites following NKX2-1 loss. A) Heatmap of TET3 occupancy at differential TET3 bound sites in KN and KNF1F2 as determined by Diffbind analysis (significance cutoff of *p*-adjusted < 0.05). B) TF motifs enriched in KN- and KNF1F2-specific TET3 bound sites. C) Mean occupancy profile of TET3 at FoxA1 bound sites in KN and KNF1F2 tumors. D) GSEA for cell type signatures on differentially bound TET3 sites identified between KN and KNF1F2 tumors. E) Fraction methylation profiles at KN- and KNF1F2-specific TET3 bound sites in K, KN, and KNF1F2 tumors. F) FoxA1 and TET3 ChIP peaks and methylation tracks at gastric pit cell genes, *Tff1* and *Gcnt3*, in KN (blue) and KNF1F2 (red) tumors.

Intersection of DEGs with DMRs revealed a strong inverse relationship between methylation and transcriptional changes. Specifically, 33.2% of genes upregulated with NKX2-1 loss were demethylated in a FoxA1/2 dependent manner (vs. 13.73% of downregulated genes; Figure 2G). This includes numerous pan gastric and pit cell marker genes such as *Hnf4a, Lgals4, Tff1, Gkn1,* and *Gcnt3*. Another 13.7% of upregulated genes were demethylated in a FoxA1/2 independent manner (vs. 5.7% of downregulated genes) including *Krt15*, a gene that is highly expressed in basal cells of the lung, esophagus, and skin. Conversely, hyperDMRs demonstrated the opposite trend, correlating more strongly with transcriptional repression. Of the 2,462 genes downregulated with *Nkx2-1* deletion, 22.8% and 7.47% gained methylation in a FoxA1/2 independent and dependent manner respectively (vs. 11.4% and 4.6% of upregulated genes). This included numerous AT2-specific marker genes such as *Sftpb*, *Dram1*, and *Cxcl15* that were methylated in a FoxA1/2 independent manner. These findings demonstrate the strong association between methylation and transcription, suggesting that methylation regulates expression of lineage-defining genes in LUAD.

Finally, we evaluated regions that gained or lost methylation specifically in KNF1F2 tumors. We identified a total of 5,262 DMRs, with 77.4% of these sites exhibiting an increase in methylation and 22.6% a decrease (Figure S4A-B, Table S2). Interestingly, while the FOX motif was the most highly enriched in KNF1F2-specific hypermethylated regions, it was not associated with hypomethylated regions (Figure S4C). Thus, FoxA1/2 appear to maintain a demethylated state at a subset of genomic loci in an NKX2-1 independent-manner. These regions also demonstrated enrichment for HNF1 motifs, TF families known to co-regulate hepatic and intestinal gene programs with FoxA1/2^36–38^. HNF1β was highly expressed across all tumor types, suggesting that FoxA1/2 may alter its activity or localization within control and KN tumors (Table S1). Gene ontology on KNF1F2-specific DMRs identified diverse cell type signatures enriched for both hyper- and hypomethylated genes (Table S4). Of note, hypomethylated genes were not significantly associated with KNF1F2 tumor cell identity (e.g., squamous or SCJ cell signatures). This may be due to the small number of DMRs within this category (n=1191) or the possibility that these gene sets are not predominantly regulated by DNA methylation. Finally, KNF1F2-specific hypermethylated regions demonstrated enrichment for fatty acid and bile acid metabolic processes, suggesting that FoxA1/2 promote demethylation of metabolism-related genes in both K and KN tumors (Table S4).

### FoxA1/2 recruit TET3 to lineage-defining gastric sites following NKX2-1 loss

Given the finding that FoxA1/2 are required for 80% of demethylation events in NKX2-1 negative LUAD, we decided to investigate the mechanism by which FoxA1/2 mediate these methylation changes. First, we performed chromatin immunoprecipitation followed by sequencing (ChIP-seq) for FoxA1 on sorted nuclei from KN tumors. This identified 17,822 FoxA1 binding sites localized primarily to promoter and distal regulatory elements (Figure S5A). Motif analysis of FoxA1 bound sites showed a strong enrichment for the FOX motif along with GI TFs seen in hypoDMRs, including KLF, HNF4, and GATA families (Figure S5B). FoxA1 bound sites significantly overlapped with genes downregulated after *Foxa1/2* deletion (Fisher’s exact test; *p*-val<10^-4^). Specifically, FoxA1 binds 56.1% of genes downregulated after deletion and 27.7% of genes upregulated, demonstrating that FoxA1/2 predominantly function as transcriptional activators in KN tumors (Figure S5C).

Given the association between FoxA1/2 binding and transcriptional activation, we next sought to determine whether FoxA1/2 directly regulate demethylation at their de novo binding sites. We found that FoxA1 bound sites were significantly more abundant in FoxA1/2 dependent-hypoDMRs as compared to independent hypoDMRs (30.3% vs. 14.2%, Fisher’s exact test; p-val<10^-4^; Figure S5D). These observations imply that FoxA1/2 directly facilitate demethylation by recruiting TET dioxygenases to their binding sites after *Nkx2-1* deletion.

To address this, we performed ChIP-seq for TET3 in KN and KNF1F2 tumors (of note, *Tet3* is expressed at similar levels in tumors of all 3 genotypes; Figure S5E). Differential peak analysis of TET3 binding identified 1100 KN-specific and 711 KNF1F2-specific sites (Figure 3A). Motif analysis of differential TET3 bound sites identified FOX, HNF4, and KLF families as the top three enriched motifs in KN samples (Figure 3B). Consistent with this, 65.9% of KN-specific TET3 peaks (n=725) directly overlapped with FoxA1 bound sites. In contrast, only 3.8% of KNF1F2-specific TET3 sites (n=27) co-localized with FoxA1. These findings suggest that FoxA1/2 directly recruit TET3 to lineage-specific sites within KN tumors. This is further supported by the loss of TET3 occupancy at FoxA1 peaks seen after *Foxa1/2* deletion (Figure 3C and S5F). Differential TET3 bound sites were then annotated to their adjacent genes for downstream functional analysis. Strikingly, KN-specific sites were significantly enriched for gastric cell types including pit and isthmus cell signatures as well as various lipid metabolic processes (Figure 3D and S5G). Enrichr cell type analysis substantiated these findings, revealing that KN-specific TET3 sites were most strongly associated with gastric and intestinal cell types (Figure S5G). Finally, we evaluated methylation signal at KN-specific TET3 sites and found a FoxA1/2 dependent reduction in methylation at TET3 peaks (Figure 3E). Inspection of specific marker genes demonstrated a clear loss of TET3 recruitment with *Foxa1/2* deletion at both pit cell (*Tff1*, *Gcnt3*, and *Gkn1*) and pan-gastric (*Hnf4a* P2 promoter, *Lgals4*, and *Ctse*) markers (Figure 3F and data not shown). Thus, FoxA1/2 are required to recruit TET3 to lineage-defining gastric sites in order to facilitate DNA demethylation.

We then evaluated KNF1F2-specific TET3 sites. Motif analysis revealed that the top enriched TF motifs were squamous lineage specifier p63 and its paralogs^39^ as well as the AP-1 complex, which is known to promote squamous differentiation in stratified epithelia^40^ (Figure 3B). GSEA and Enrichr cell type analysis on KNF1F2-specifc TET3 sites showed a significant association with basal cell and squamous signatures (Figure 3D and S5G). KNF1F2-specific TET3 binding was identified at squamous epithelial markers *Krt5* and *Krt14* as well as genes highly expressed in the SCJ of the stomach including *Chil4*, *Mmp7*, *Fbln2*, and *Rbp7* (Figure S5J and data not shown). These findings are consistent with our current and previously published datasets showing that KNF1F2 tumors adopt a squamous or SCJ-like state ^30^. Intriguingly, when we evaluated methylation signal at KNF1F2-specific TET3 sites, we saw no significant difference in the fraction methylation between tumor types. Instead, these regions were lowly methylated in all tumor samples, suggesting that KNF1F2-specific TET3 sites are demethylated during either lung development or upon KRAS^G12D^ activation. These findings suggest that methylation is not a major mechanism regulating transcription of these genes. It also raises the possibility that TET3 may perform non-canonical functions at KNF1F2-specific sites to regulate the epigenetic state (e.g., recruitment of chromatin-modifying enzymes)^41^. Overall, these data show that FoxA1/2 loss results in de novo TET3 binding to squamous-specific sites that are stably hypomethylated, regardless of TET3 localization

### FoxA1/2 are required for H3K27ac deposition and enhancer-promoter interactions at gastric-specific sites in NKX2-1-negative LUAD

In addition to modifying DNA methylation, FoxA1/2 are known to direct histone PTMs via recruitment of histone-modifying enzymes^23^. We have previously shown that de novo FoxA1/2 bindings sites exhibit an increase in H3K27ac, a marker of active regulatory elements, following NKX2-1 loss^18^. However, it is unknown whether these epigenetic changes are dependent on FoxA1/2 or whether they coincide with lineage-specific TET3 binding and DNA demethylation.

To answer this, we performed ChIP-seq for H3K27ac on sorted nuclei from KN and KNF1F2 tumors. Differential peak analysis identified 4,092 KN-specific and 4,217 KNF1F2-specific H3K27ac sites (Figure 4A). Consistent with our TET3 motif analysis, we found that KN-specific H3K27ac sites were significantly enriched for FOX and HNF4 motifs (Figure 4B). Accordingly, 51.8% of KN-specific H3K27ac sites (n=2119) directly overlapped with FoxA1 bound sites (versus 5.1% of KNF1F2-specific H3K27ac sites). H3K27ac occupancy also demonstrated a significant depletion at FoxA1 peaks following *Foxa1/2* deletion (Figure S6A-B). Cell type enrichment analysis revealed that KN-specific sites were most significantly associated with gastric pit and goblet cell signatures, showing that FoxA1/2-dependent H3K27ac sites occur at gastric-defining genes (Figure S6C). This was substantiated by Enrichr analysis showing that KN-specific sites were most strongly associated with human stomach tissue (Table S3). Finally, we found that KN-specific HK27ac sites were enriched for mucin biosynthesis and numerous metabolic processes (Figure S6D). Thus, FoxA1/2 are required for not only TET3 recruitment, but H3K27ac deposition at genomic sites involved in regulation of gastric identity and cell metabolism.

**Figure 4:**
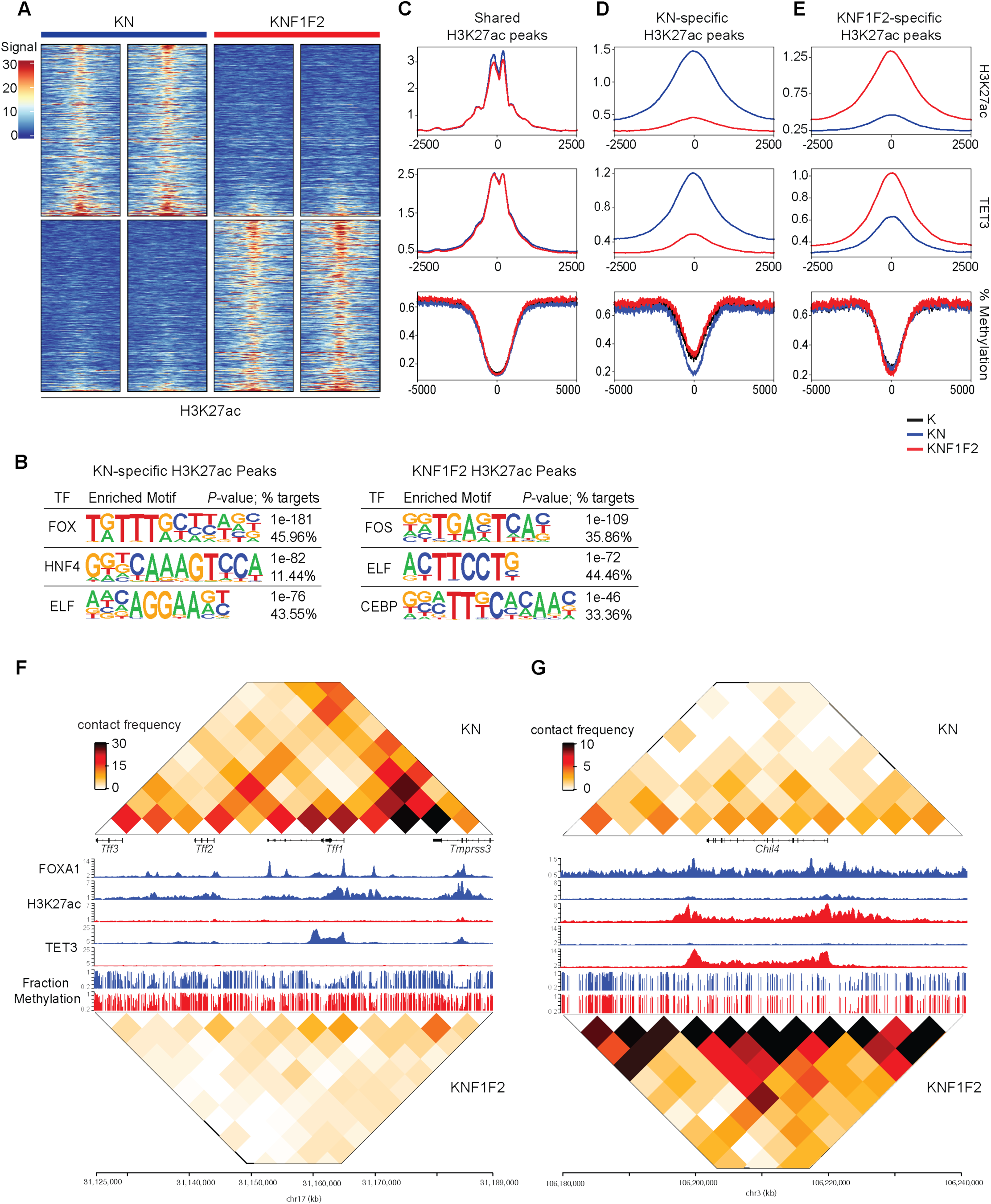
FoxA1/2 are required for H3K27ac deposition and E-P interactions at gastric-specific sites in NKX2-1-negative LUAD. A) Heatmap of H3K27ac occupancy at differential H3K27ac sites in KN and KNF1F2 as determined by Diffbind analysis (significance cutoff of *p*-adjusted < 0.05). B) TF motifs enriched in KN- and KNF1F2-specific H3K27ac sites. C) H3K27ac and TET3 ChIP occupancy and fraction methylation profiles at shared H3K27ac sites in K, KN, and KNF1F2 tumors. D) H3K27ac and TET3 ChIP occupancy and fraction methylation profiles at KN-specific H3K27ac sites in K, KN, and KNF1F2 tumors. E) H3K27ac and TET3 ChIP occupancy and fraction methylation profiles at KNF1F2-specific H3K27ac sites in K, KN, and KNF1F2 tumors. F) Heatmap of H3K27ac contact frequency with FoxA1, TET3, and H3K27ac ChIP peaks and methylation tracks at genomic loci containing gastric genes, *Tff1, Tff2,* and *Tff3*, in KN (blue) and KNF1F2 (red) tumors. G) Heatmap of H3K27ac contact frequency with FoxA1, TET3, and H3K27ac ChIP peaks and methylation tracks at SCJ marker, *Chil4*, in KN (blue) and KNF1F2 (red) tumors.

Next, we evaluated KNF1F2-specific H3K27ac sites. The most highly enriched TF motif for these regions was bound by the AP1 family, which can regulate squamous differentiation^40^ and was also seen in KNF1F2-specific TET3 peaks (Figure 3B). Although GSEA did not identify enrichment of a specific identity program, Enrichr analysis with the GTEx database showed that KNF1F2-specific H3K27ac sites were most significantly associated with esophagus (Figure S6E). Moreover, we saw significant H3K27ac accumulation at markers of squamous identity, including *Trp63* and *Krt5,* as well as SCJ genes, *Chil4* and *Mmp7*, in KNF1F2 tumors (Figure S6F). TF enrichment analysis using Encode and ChEA ChIP-X datasets showed a significant enrichment for SOX2 and TP63 (Table S3). Thus, a portion of the differential H3K27ac sites occupy lineage-defining squamous and SCJ sites in KNF1F2 tumors. Altogether, these results show that FoxA1/2 are required for H3K27ac deposition at gastric-specific sites and that, in the absence of FoxA1/2, H3K27ac accumulates at genes relating to squamous differentiation.

Given the association between H3K27ac deposition and tumor cell lineage, we next wanted to determine whether differential H3K27ac sites coincided with altered TET3 recruitment and DNA methylation patterns. As such, we looked at the distribution of TET3 ChIP-seq signal and fraction methylation at KN-specific, KNF1F2-specific, and shared H3K27ac peaks in KN and KNF1F2 tumors. Although shared H3K27ac peaks exhibited similar levels of TET3 binding and DNA methylation (Figure 4C), KN-specific H3K27ac sites demonstrated significantly higher TET3 recruitment and lower DNA methylation in KN tumors (Figure 4D). These data highlight the genomic colocalization of FoxA1/2 dependent epigenetic changes and show that lineage-specific H3K27ac sites undergo FoxA1/2-mediated TET3 recruitment and DNA demethylation. In contrast, KNF1F2-specific H3K27ac sites exhibited increased TET3 localization with no methylation changes relative to K or KN tumors (Figure 4E). These results align with the aforementioned finding that KNF1F2-specific TET3 sites do not coincide with differential methylation and imply that alternative epigenetic mechanisms regulate transcription of these sites.

As many of the affected H3K27ac sites are distant to gene promoters, they likely regulate their target genes through enhancer-promoter (E-P) loops. We therefore performed H3K27ac HiChIP on sorted nuclei from KN and KNF1F2 tumors, which identified 473,024 and 712,989 significant interactions between H3K27ac peaks in KN and KNF1F2 tumors respectively. Next, we categorized H3K27ac contacts by the presence of only one transcription start site (TSS) at one anchor to identify E-P loops specifically. We then assessed the global changes of FoxA1/2 dependent cis-regulatory E-P interactions caused by FoxA1/2 depletion. As expected, shared H3K27ac sites present in both tumor types exhibited no significant difference in the number of E-P contacts (Figure S7A). However, KN-specific H3K27ac sites demonstrated a significant reduction in E-P loop interactions following *Foxa1/2* deletion, showing that FoxA1/2 are required for cis-regulatory interactions at these H3K27ac sites. Conversely, KNF1F2-specific H3K27ac sites demonstrated a significant increase in E-P interactions with *Foxa1/2* deletion. Intersection of E-P loops with DEGs identified after FoxA1/2 loss show that genes upregulated in KN or KNF1F2 tumors exhibit significantly more E-P loops within each respective tumor type (Figure S7B). These findings suggest that genes activated in KNF1F2 tumors are not epigenetically regulated by methylation (given the lack of demethylation that occurs with FoxA1/2 loss) but by the presence of active histone modifications and cis-regulatory interactions.

Lastly, we examined the epigenetic landscape at selected lineage-defining gastric and SCJ marker genes. Examination of the genomic region containing GI genes *Tff1*, *Tff2*, and *Tff3* showed numerous sites bound by FoxA1 in KN tumors (Figure 4F). These FoxA1 bound sites directly overlapped with regions demonstrating FoxA1/2 dependent demethylation, TET3 recruitment, H3K27ac deposition, and E-P looping. In particular, we see strong physical interactions between an enhancer located approximately 20 kb upstream from the *Tff1* TSS and the promoters of both *Tff1* and *Tff2*, which is disrupted after *Foxa1/2* deletion. Similar FoxA1/2 dependent epigenetic changes were seen in the genomic loci containing gastric pit cell markers, *Gcnt3* and *Gkn1*, as well as the pan-gastric marker *Hnf4a* (Figure S7C and data not shown). Thus, FoxA1/2 are required to comprehensively reprogram the epigenetic landscape of lineage-defining genes to promote a gastric differentiation state in NKX2-1-negative LUAD.

Evaluation of SCJ marker genes highly expressed in KNF1F2 tumors revealed contrasting epigenetic patterns. For example, SCJ marker *Chil4* does not contain any FoxA1 binding sites nor does it exhibit an active chromatin state in KN tumors (Figure 4G). However, following *Foxa1/2* deletion, we see a significant accumulation of H3K27ac, TET3 binding, and local E-P contacts. Interestingly, while there is a slight reduction in methylation at the *Chil4* promoter, it does not meet the significance threshold as this site has similar methylation levels in both KN and KNF1F2 tumors. This data supports our prior conclusions that most genes exhibiting increased TET3 recruitment after *Foxa1/2* deletion are not differentially methylation. Examination of another SCJ marker, *Mmp7*, demonstrates a similar trend in which H3K27ac deposition, TET3 binding, and E-P looping occur only after FoxA1/2 loss (Figure S7D). Notably, the promoter of *Mmp7* does contain a small KNF1F2-specific hypoDMR, suggesting that a small subset of SCJ marker genes may be regulated by methylation.

### FoxA1/2 are required to maintain gastric identity, but not hypomethylation of gastric regulatory elements, in NKX2-1-negative LUAD

DNA methylation is a central epigenetic mark that can be stably inherited by progeny cells in order to maintain cellular identity. As such, mechanisms required to establish methylation patterns may not be necessary for their maintenance. For example, FoxA1/2 are required to establish hypomethylated patterns at liver-specific enhancers *in vivo*, but they are dispensable for their maintenance^24^. Thus, we decided to investigate whether FoxA1/2 are required to maintain gastric gene expression and hypomethylated patterns at gastric loci following *Nkx2-1* deletion in LUAD.

To address this question, we employed an *in vitro* 3D organoid system that uncouples *Foxa1/2* deletion from NKX2-1 loss. After deriving organoids from KNF1F2 tumors, we noticed that *Nkx2-1* is often stochastically downregulated. This uncoupling event likely occurs due to the growth advantage seen in KRAS^G12D^-driven lung neoplasia following *Nkx2-1* repression^18,42^. Thus, we derived and screened multiple KNF1F2 organoid lines to identify those that had (1) uniformly downregulated *Nkx2-1* expression, (2) retained *Foxa1/2* expression, and (3) adopted a gastric state. This led to the selection of two organoid lines, KG1A and 22E, that met all criteria. We then confirmed that treatment with 4-hydroxytamoxifen (4-OHT) *in vitro* successfully deleted *Foxa1/2* and resulted in downregulation of the canonical FoxA1/2 target gene, *Hnf4a* (Figure 5A).

**Figure 5:**
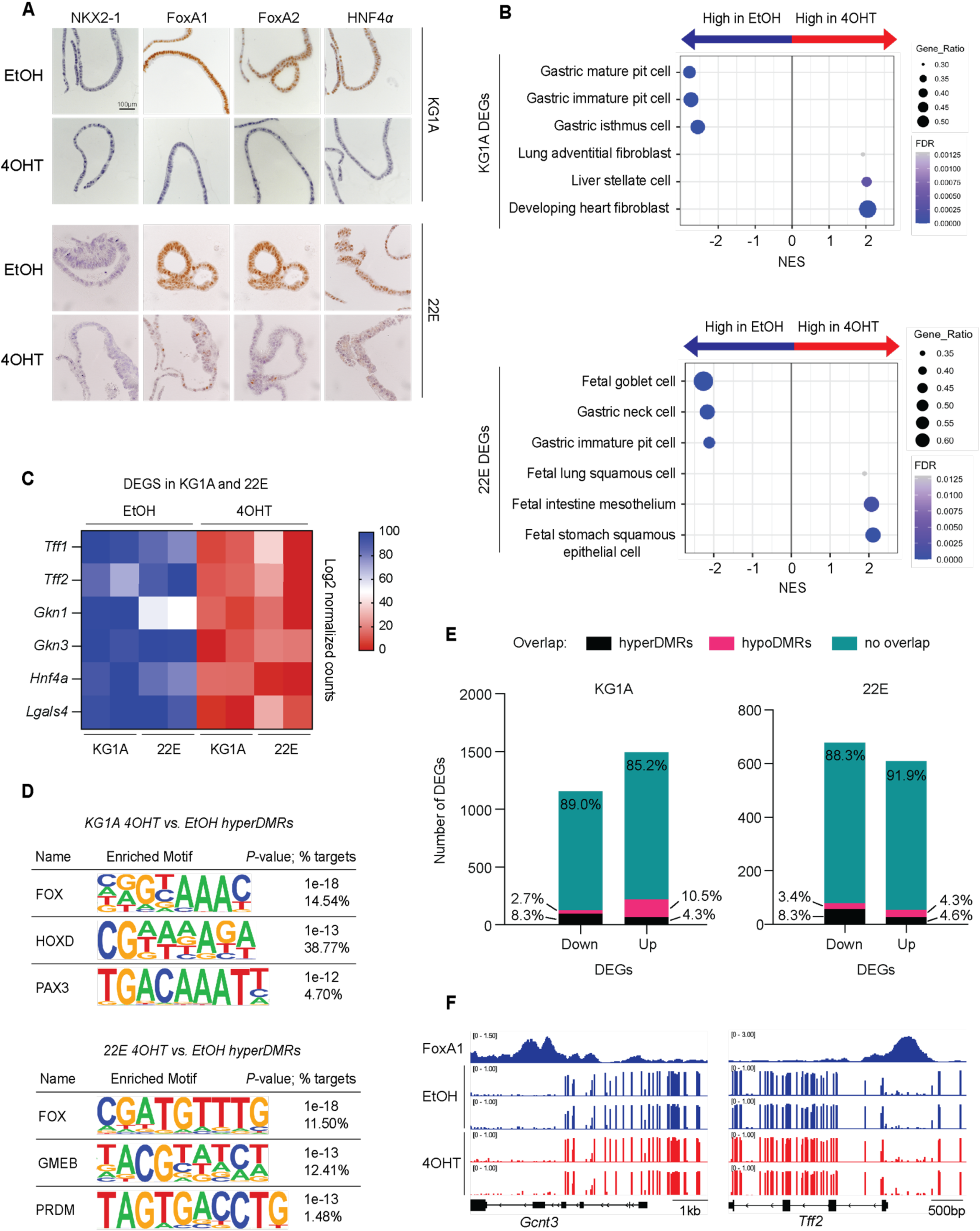
FoxA1/2 are required to maintain gastric identity, but not hypomethylation of gastric regulatory elements, in NKX2-1-negative LUAD. A) Representative images of KG1A and 22E organoids two weeks after treatment with EtOH or 4-OHT. IHC of NKX2-1, FoxA1, FoxA2, and HNF4α shown. B) GSEA for cell type signatures on DEGs identified in EtOH vs. 4OHT for KG1A and 22E organoids. Organoids collected two weeks post treatment. C) Heatmap showing log2 normalized counts of GI genes in EtOH vs. 4OHT for KG1A and 22E organoids. D) TF motifs enriched in hyperDMRs identified in EtOH vs. 4OHT for KG1A and 22E organoids. E) Intersection of DEGs and DMRs identified in EtOH vs. 4OHT for KG1A and 22E organoids. Less than 10% downregulated genes show a corresponding increase in methylation. F) FoxA1 ChIP peaks and methylation tracks at gastric genes, *Gcnt3* and *Tff2*, in KG1A organoids treated with EtOH (blue) or 4OHT (red).

To determine whether FoxA1/2 are required for maintenance of a gastric identity, we performed bulk RNA-seq on KG1A and 22E organoids in the presence and absence of FoxA1/2. We found that deletion of *Foxa1/2 in vitro* led to extensive transcriptional changes with 2,144 DEGs identified in KG1A and 1,286 DEGs in 22E (log_2_FC > 1; padj < 0.05, Figure S8A, Table S5). Intersection of these datasets with our *in vivo* RNA-seq revealed a considerable overlap, whereby 28.6% and 20.3% of the genes downregulated *in vivo* with FoxA1/2 loss were also downregulated in KG1A and 22E respectively (Figure S8B). GSEA analysis of cell identity programs closely mirrored results of *Foxa1/2* deletion *in vivo* (Figure 5B). Both organoid lines showed a strong enrichment for gastric cell types, particularly pit and goblet cell signatures, that were lost upon *Foxa1/2* deletion. Individual gastric targets including pit cell markers, *Tff1* and *Gkn1*, and mucous neck cell markers, *Tff2* and *Gkn3,* were substantially downregulated with FoxA1/2 loss in both organoid lines (Figure 5C). Pathway analysis also identified FoxA1/2-dependent metabolic processes in vitro (Figure S8C) that were similar to those observed in vivo (Figure S3D). On the other hand, *Foxa1/2* deleted lines showed enrichment for multiple cell type signatures and inflammatory processes. Of note, KG1A and 22E were significantly associated with distinct identities after deletion, which partially recapitulates cell fate heterogeneity seen after *Foxa1/2* deletion *in vivo*. Overall, these data demonstrate that FoxA1/2 are required for maintaining a gastric differentiation state in NKX2-1-negative LUAD.

We next sought to determine whether FoxA1/2 are required for maintenance of DNA hypomethylation patterns at lineage-defining genes. We performed whole genome EM-seq on KG1A and 22E in the presence and absence of FoxA1/2 as well as ChIP-seq for FoxA1 in both lines. Interestingly, loss of FoxA1/2 in NKX2-1-negative organoids led to relatively modest methylation changes when compared with the results of concomitant *Foxa1/2;Nkx2-1* deletion *in vivo* (Figure 2). In KG1A and 22E, we identified a total of 3,914 and 2,280 DMRs respectively (Figure S8D). FoxA1 ChIP-seq identified 21,827 peaks in KG1A and 26,091 in 22E, localized primarily to intronic and intergenic regions (i.e., potential enhancer sites, Figure S8E). To determine whether FoxA1/2 directly alter methylation patterns, we intersected FoxA1 bound sites with DMRs. Regions that gained methylation after FoxA1/2 loss (“hyperDMRs”) demonstrated a significant overlap with FoxA1 bound sites (as compared with regions that lost methylation or “hypoDMRs”). For example, in KG1A, 27.6% of hyperDMRs directly overlapped with FoxA1 bound sites versus 6.24% of hypoDMRs (Fisher’s exact test; p-val<10^-4^). These findings were substantiated by motif analysis which showed a significant enrichment for the FOX motif in hyperDMRs, but not hypoDMRs, for both organoid lines. These data demonstrate the role of FoxA1/2 in maintaining a demethylated state at target sites in KN organoids.

Given the association between FoxA1 binding and demethylation, we next investigated whether FoxA1/2 are responsible for maintaining hypomethylated patterns at gastric-specific sites. First, we performed cell type enrichment analysis on hyperDMRs, which revealed an enrichment for various midbrain signatures but no GI cell types (Table S4). Although motif analysis of hyperDMRs showed a strong enrichment for the FOX motif, it lacked other GI TF’s including HNF4 and KLF families that were found *in vivo* (Figure 5D). Intersection of DMRs with transcriptional changes revealed a minor overlap: less than 10% of genes downregulated with *Foxa1/2* deletion had a corresponding increase in methylation for both lines (Figure 5E). In fact, numerous gastric marker genes both bound by FoxA1 and downregulated with *Foxa1/2* deletion, exhibited no change in methylation. For example, gastric targets, *Tff2* and *Gcnt3*, which were significantly downregulated with FoxA1/2 loss showed no increase in DNA methylation (Figure 5F). While most gastric targets did not undergo substantial methylation changes, there were several notable exceptions including *Tff1* and *Gkn3,* which were significantly methylated upon FoxA1/2 loss (Figure S8F).

Altogether, these findings demonstrate the diverging roles of FoxA1/2 in regulation of gastric gene transcription and methylation. Although FoxA1/2 are required to maintain a gastric cellular identity in NKX2-1-negative LUAD, they are not required to maintain hypomethylated patterns at most gastric loci. These findings substantiate previous work^8,24^ showing that lineage-specific DNA methylation patterns are stable and do not always require the TFs with which they were established.

### Oncogenic KRAS is required for FoxA1/2-dependent transcriptional activation and demethylation of gastric pit cell genes in NKX2-1-negative LUAD

Aberrant KRAS signaling plays a pivotal role in LUAD pathogenesis, inducing tumor cell survival and proliferation via activation of its canonical target, extracellular signal-related kinase (ERK). The importance of ERK activity is underscored by the observation that small molecule KRAS inhibitors block ERK activation to a greater extent than other downstream signaling pathways^43,44^. In addition to its roles in tumor progression, ERK signaling is known to regulate cell differentiation and development^45^. In mouse embryonic stem cells, ERK signaling suppresses pluripotent gene expression to induce an endodermal cell state^46,47^. Perturbations in ERK signaling are also associated with developmental abnormalities in tissues including the bone, heart, and brain^45^. We have previously shown that RAF/MEK/ERK activity modulates the exact gastric identity adopted by NKX2-1-negative LUAD. Pharmacological inhibition of RAF/MEK in a BRAF^V600E^ model of IMA represses the gastric pit cell program while inducing markers of gastric chief and tuft cells^4^. We have also shown that pulmonary *Nkx2-1* deletion in the absence of KRAS^G12D^ induces small mucinous alveolar hyperplasias that express pan-gastric markers such as HNF4α, but not the gastric pit cell marker GKN1^18^ by immunostaining.

Given these findings, we decided to profile the transcriptional changes that occur with NKX2-1 loss in normal lung alveolar cells to determine the effect of oncogenic KRAS on tumor cell identity. Bulk RNA-seq demonstrated that cells derived from *Nkx2-1*-deleted hyperplasias assumed a distinct transcriptional state from their KN counterparts (n=2,465 DEGs identified, log_2_FC>1; padj<0.05; Figure S9A; Table S1). *Dusp6*, a marker of ERK activity, was significantly upregulated in KN tumors compared to N, consistent with increased RAF/MEK activity downstream of KRAS^G12D^. GO analysis demonstrated a significant enrichment for gastric pit cell signatures in KN tumors (Figure 6A), as exemplified by genes such as *Tff1*, *Gkn1*, and *Gcnt3* (Figure 6B). Given the fact that both KN tumors and *Nkx2-1*-deleted hyperplasias upregulate pan-gastric markers, we evaluated the extent to which each sample is associated with distinct gastric identities. Using single sample GSEA (ssGSEA), we found that N hyperplasias are also enriched for pit cell signatures relative to K and KNF1F2 tumors, but to a much lesser extent than KN tumors (Figure 6C and S9B). Other gastric programs including tuft cell signatures demonstrated similarly high enrichment in KN and N samples compared with K and KNF1F2 (Figure S9C). Consistent with this, pan gastric markers *Hnf4a* and *Ctse* were expressed at similarly high levels in KN and N samples (Figure S9D). Altogether, these results show that *Nkx2-1* deletion within AT2 cells is sufficient to activate elements of a gastric differentiation program, but it is the combination of NKX2-1 loss with oncogenic KRAS signaling that is required for maximum induction of a gastric pit cell identity.

**Figure 6:**
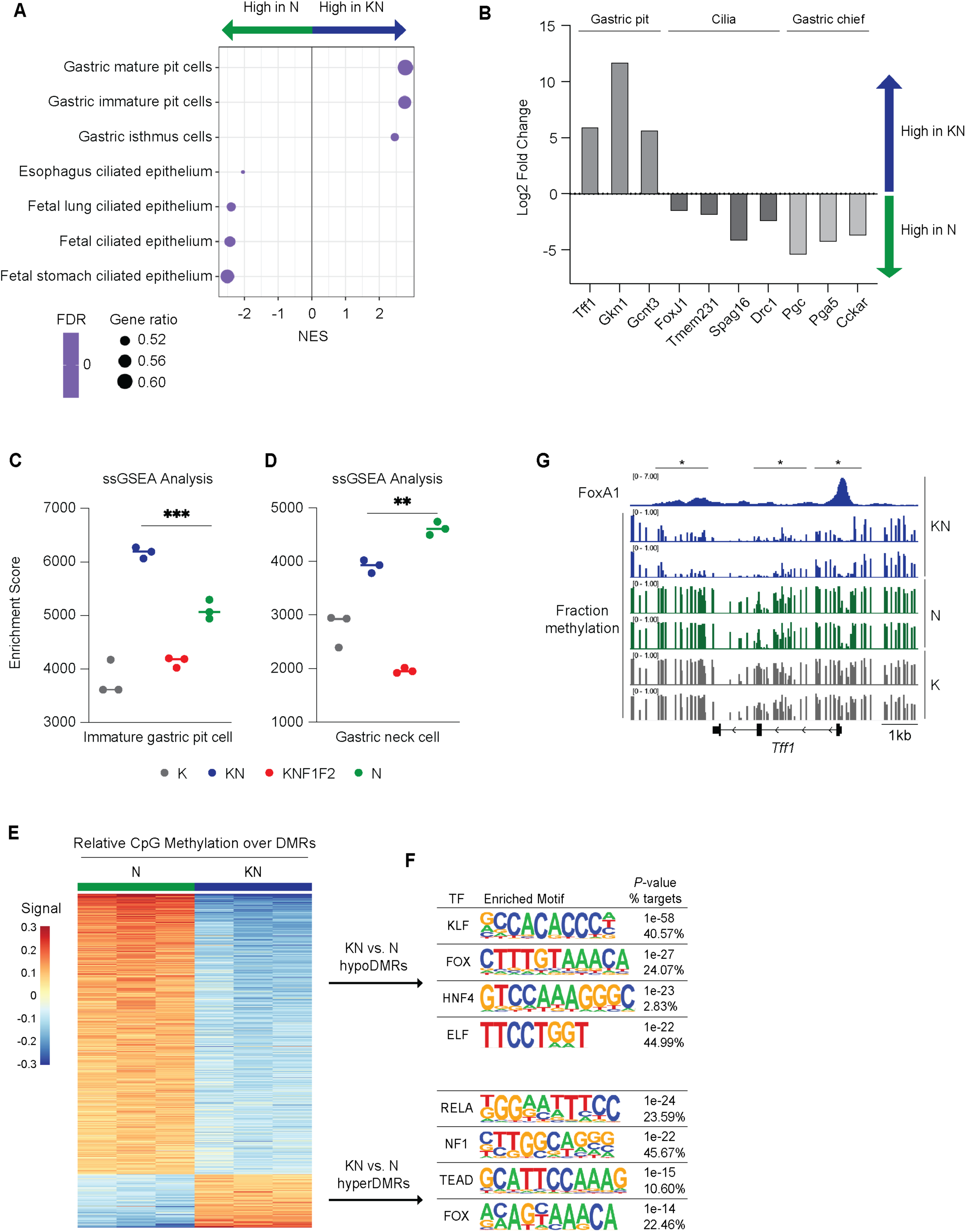
Oncogenic KRASG12D is required for transcriptional activation and demethylation of gastric pit cell genes in NKX2-1-negative LUAD. A) GSEA for cell type signatures on DEGs identified in KN vs. N samples. B) Log2 fold change values of genes comprising the gastric pit, gastric chief, or ciliated cell signatures in KN vs. N tumors. C) Enrichment score determined by ssGSEA for immature gastric pit cell signature in K, KN, KNF1F2, and N samples. \*\*\**P*-value < 10^-3^ in KN vs. N cells, t-test. D) Enrichment score determined by ssGSEA for gastric neck cell signature in K, KN, KNF1F2, and N samples. \*\**P*-value < 10^-2^ in N vs. KN cells, t-test. E) Heatmap of mean relative CpG methylation over DMRs identified in KN vs. N samples. Samples collected 14-weeks post tumor initiation. F) TF motifs enriched in KN vs. N DMRs. G) FoxA1 ChIP peaks and methylation tracks at gastric pit cell gene, *Tff1*, in KN, N, and K samples.

In contrast to KN tumor cells, N cells demonstrated a strong enrichment for fetal ciliated epithelial cell types (Figure 6A). GO and Enrichr analysis corroborated these findings, showing a significant association with cilia and microtubule gene sets (Figure S9E and Table S3). ChIP-X Enrichment Analysis 3 (ChEA3) further supported these results, showing that FoxJ1, a master regulator of ciliated cell differentiation, was one of the top two TFs enriched in N cells (Figure S9F). Specific genes involved in cilia regulation and assembly were strongly upregulated in N lesions including *Foxj1*, *Tmem231*, *Spag16*, and *Drc3* (Figure 6B). In addition to cilia-related gene sets, N cells also upregulated gastric chief cell markers including *Pgc*, *Pga5*, and *Cckar* (Figure 6B) and exhibited a strong association with gastric neck cell signatures (Figure 6D). Of note, stomach chief cell signatures were associated with N samples but did not meet the significance *p*-value threshold of 5%. Interestingly, the cell type enrichment profiles for KN and N samples appear to reflect the anatomic distribution of gastric glands with higher levels of MAPK signaling (e.g., KN tumors) activating a surface pit cell state while lower levels (e.g., N cells) activate more internal cell fates including neck and chief cell identities. Overall, these data expand upon our previous findings to show that KRAS^G12D^ is required for full activation of a gastric pit cell state in NKX2-1-negative LUAD and that, in the absence of a driver oncogene, NKX2-1-negative lung epithelial cells adopt a ciliated epithelial state with co-expression of a subset of gastric chief and neck cell markers.

We next used whole genome EM methyl-seq on sorted nuclei to determine the impact of KRAS^G12D^ on DNA in NKX2-1 negative lung cells. We identified 10,620 DMRs between N and KN nuclei, with 9,026 sites exhibiting increased methylation compared to KN samples and 1,594 sites exhibiting a decrease (Figure 6E, Table S2). HOMER analysis identified the FOX motif in both N and KN DMRs (Figure 6F). However, N samples lacked enrichment for additional GI TF motifs present in KN samples including KLF and HNF4 families; instead, they were associated with an assorted group of motifs including RELA, NF1, and TEAD. Intriguingly, KN samples did not exhibit higher expression of *Foxa1/2* or *Hnf4a/g*, suggesting that KRAS^G12D^ might alter their function through post-translational mechanisms such as phosphorylation and/or changes in the composition of TF complexes. To determine the role of oncogenic KRAS on FoxA1/2-dependent demethylation events (Figure 2A, purple), we intersected both datasets and found that 29.8% of all FoxA1/2-dependent hypoDMRs (n=2,481) directly overlapped with KRAS-dependent DMRs. 26.9% of these FoxA1/2-KRAS co-dependent sites (n=667) were directly bound by FoxA1, and 72.4% (n=1797) are within 10 kb of a FoxA1 binding site. Moreover, 13.8% of KN-specific hypoDMRs contained a FoxA1 binding site (versus 4.6% of N-specific hypoDMRs; Fisher’s exact test, *p*-val<10^-4^). These data demonstrate a close association between FoxA1 binding and KRAS-dependent DMRs, and also raise the question of whether KRAS^G12D^ expression promotes FoxA1/2 binding to these sites to regulate cell identity.

GO analysis on DMGs showed that the top two cell type signatures enriched for KN hypomethylated genes were gastric immature and mature pit cells (Figure S9G). Individual pit cell markers *Tff1*, *Gkn1*, and *Gcnt3* as well as gastric targets *Lgals4* and *Anxa8* all demonstrated demethylation that was dependent on the combination of KRAS^G12D^ expression and *Nkx2-1* deletion (Figure 6G and S9H). Inspection of key KRAS-dependent pit cell markers showed that most genes either directly overlapped with FoxA1 bound sites (e.g., *Tff1*) or contained local FoxA1 binding sites (e.g., *Gcnt3* and *Gkn1*). In contrast, pan gastric markers including *Hnf4a* and *Ctse* were demethylated upon NKX2-1 loss independent of KRAS^G12D^ activation. Evaluation of N-specific hypomethylated genes did not reveal any significant associations with specific cell types or gene ontology signatures (perhaps due to the small number of sites identified). Moreover, cilia marker genes *Foxj1*, *Tmem231*, and *Drc3* did not undergo demethylation despite being upregulated in N hyperplasias, suggesting that transcriptional regulation of these genes is methylation independent. Altogether, these findings show that KRAS signaling is essential for demethylation of select gastric and pit cell marker genes. Thus, it is the combined action of oncogenic KRAS and FoxA1/2 that leads to demethylation and transcriptional activation of specific gastric loci during the lineage switch that follows the loss of NKX2-1 in LUAD.

## DISCUSSION

Cell plasticity is intricately tied to the evolutionary fitness of a cancer cell. The ability of tumor cells to alter their identity in response to selective pressures (natural or therapy-induced) is essential for their continued growth, survival, and malignant potential. Therefore, it is imperative to define the precise molecular mechanisms, including the transcriptional and epigenetic factors, that control cancer cell identity. Here, we show that pioneer factors FoxA1/2 reprogram the epigenetic landscape of NKX2-1-negative LUAD to facilitate cancer cell lineage switching. Following *Nkx2-1* deletion, FoxA1/2 localize to regulatory elements of gastric-specific sites where they coordinate DNA demethylation, H3K27ac deposition, and E-P interactions (Figure 7). We also show that FoxA1/2 recruit the TET3 dioxygenase to lineage-defining genes, providing a likely mechanism for their demethylation.

**Figure 7:**
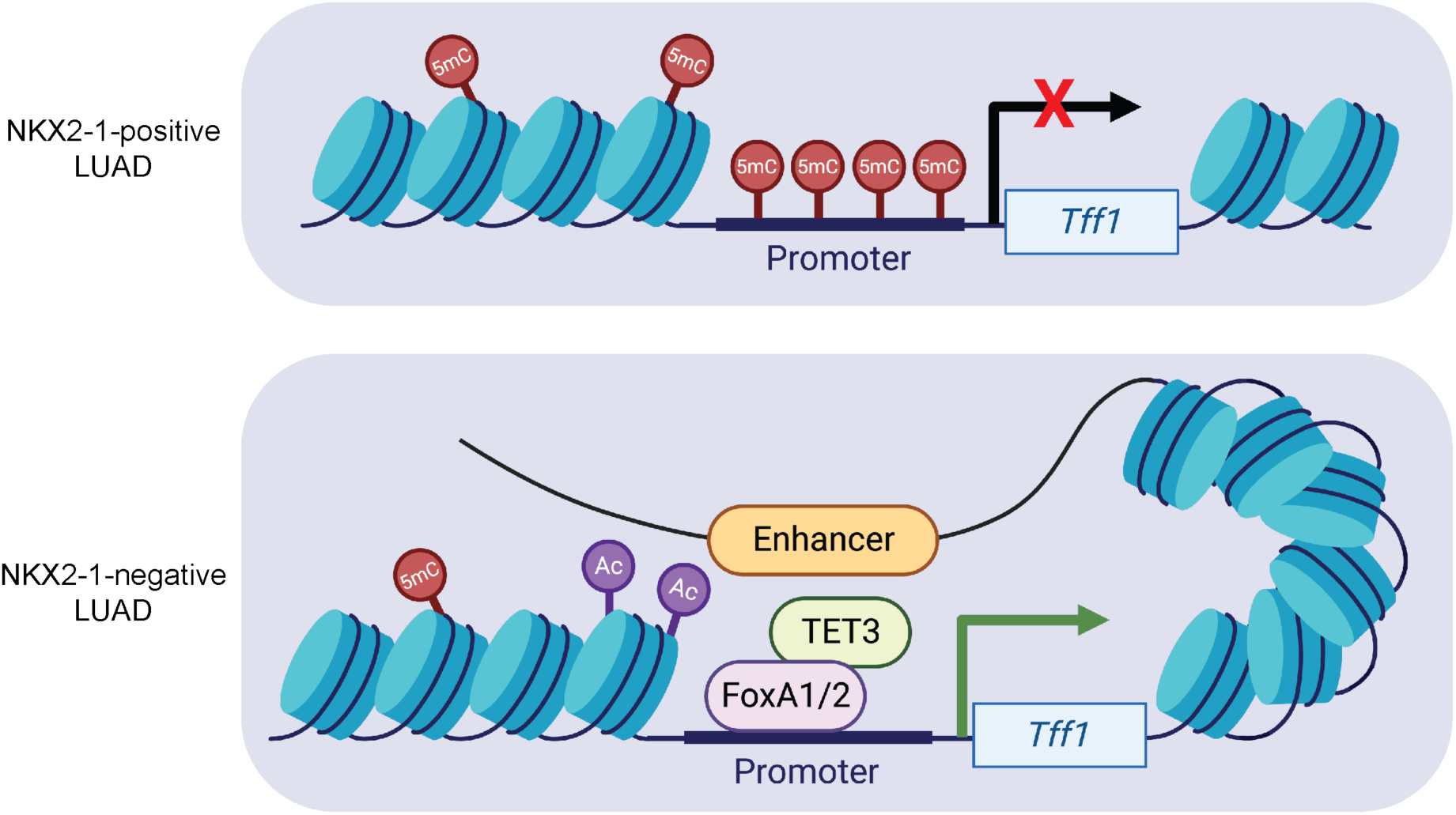
FoxA1/2 reprogram the epigenetic landscape of NKX2-1-negative LUAD to drive cancer cell lineage switching. A) Working model of FoxA1/2-dependent epigenetic changes that drive the pulmonary-to-gastric lineage switch in LUAD after NKX2-1 loss.

The systematic rewiring of the LUAD epigenetic state to promote a gastric identity is likely linked to the developmental origins of the lungs. Foregut endoderm, a portion of the gut tub that forms numerous GI structures, also gives rise to the primitive lung. As endodermal lineage specifiers, FoxA1/2 regulate the development and differentiation of both pulmonary and GI tissues derived from foregut endoderm. Therefore, deletion of the pulmonary lineage specifier *Nkx2-1* likely releases FoxA1/2 from their roles in maintaining a pulmonary cell fate and allows them to adopt alternative functions in foregut differentiation (i.e., structuring the epigenetic landscape to facilitate a gastric identity). Indeed, our findings support a model where FoxA1/2 bind gastric targets after *Nkx2-1* deletion and directly recruit enzymes that modify DNA methylation and histone PTMs to facilitate epigenetic alterations. In other endodermal tissues such as the liver, FoxA1/2 are known to mediate similar epigenetic changes including demethylation of lineage-specific enhancers and maintenance of an active chromatin state^24,48^. However, the specific epigenetic functions performed by FoxA1/2 during stomach differentiation are not as well understood^49^. Thus, our findings may provide insight into how FoxA1/2 normally regulate formation of gastric cell fate during endodermal development.

MAPK signaling is known to regulate GI epithelial differentiation^50^. Here, we show that oncogenic signaling downstream of KRAS^G12D^ alters the precise differentiation state adopted by NKX2-1-negative LUAD tumors cells. Specifically, KRAS^G12D^ activates a gastric pit cell program, whereas *Nkx2-1* deleted, KRAS wild type hyperplasias express transcriptional programs of ciliated cells and gastric chief cells. Based on our prior work^4^, these effects are likely mediated by RAF/MEK signaling downstream of KRAS^G12D^. These findings align with recent studies performed in human gastric organoids showing that MAPK signaling activation is essential for adoption of a pit cell program but detrimental for chief or parietal cell states^51^. Interestingly, we find that regardless of MAPK signaling activation, the FoxA motif is associated with sample-specific unmethylated regions. This poses the fundamental question of how MAPK signaling alters FoxA1/2 activity to modulate cell identity, suggesting that FoxA1/2 genomic localization or co-factor interactions may be modified by RAF/MEK activity. Additional studies will be needed to elucidate the precise mechanism by which MAPK signaling modulates FoxA1/2 function. Results from these studies also have implications for understanding the cell identity that LUAD adopts following targeted KRAS inhibition and the role of FoxA1/2 in facilitating this differentiation program.

Following *Nkx2-1* deletion, we saw robust TET3 recruitment and demethylation at lineage-defining gastric pit cell genes. However, in FoxA1/2-negative tumors, TET3 recruitment to squamous and SCJ genes did not correspond with reduced methylation. Instead, these sites exhibited hypomethylated patterns in all tumor types regardless of TET3 binding. This suggests that genomic regions associated with squamous and SCJ differentiation are demethylated during lung development or following KRAS^G12D^ activation and that these patterns are stably propagated within tumor cells. These data raise the question of whether increased TET3 localization to these sites following *Foxa1/2* deletion has a functional consequence. Recent studies have shown that TET enzymes directly regulate H3K27 modification via recruitment of histone-modifying enzymes that is independent of its catalytic activity^41^. Thus, TET3 may be involved in modulating histone PTMs and chromatin dynamics at lineage-specific sites. Studies aimed at understanding the role of TET3 and its co-factors in KNF1F2 tumor differentiation are highly relevant given the fact that squamous transformation occurs during both LUAD tumor progression (e.g., human adenosquamous carcinomas^52^) and as a mechanism of acquired resistance to KRAS^53^ and EGFR^54^ inhibition. Given the genomic overlap between TET3 binding and H3K27ac deposition within KN tumors, it would also be interesting to see whether TET3 acts in concert with FoxA1/2 to regulate the histone landscape of LUAD.

This study examined the molecular mechanisms regulating epigenetic reprogramming during cancer cell lineage switching. Future experiments are needed to delineate the precise role of epigenetic alterations in terms of gene transcription (i.e., determine which mechanisms regulate transcription versus mark active chromatin). Additionally, it remains to be determined whether specific epigenetic changes occur in a specific temporal sequence. For example, is demethylation required for H3K27ac deposition and E-P looping at lineage-specific genes? This knowledge will be imperative for the development of therapeutic strategies aimed at preventing or counteracting identity-specific vulnerabilities.

## Acknowledgements

We are grateful to members of the Snyder lab for suggestions and comments. We thank Brian Dalley for sequencing expertise, Jay Gertz for ChIP-seq expertise, James Marvin for FACS expertise, and Ian MacIsaac for assistance with data analysis. The results published here are in part based upon data generated by the TCGA Research Network: https://www.cancer.gov/tcga. E.L.S. was supported by grants from the NIH (R01CA212415, R01CA240317 and R01CA237404), the American Lung Association (LCD-821670), and institutional funds (Department of Pathology and Huntsman Cancer Institute/Huntsman Cancer Foundation, University of Utah). K.G. and G.F. were supported by the NIH/NCI (F31CA275266 and F31CA275328). G.F. was also supported by Genetics Training Grant (T32GM141848). Research reported in this publication utilized shared resources (including High Throughput Genomics, Bioinformatics, Flow Cytometry, and Biorepository and Molecular Pathology) at the University of Utah and was supported by the National Cancer Institute of the National Institutes of Health under award number P30CA042014. Work in the flow cytometry core was also supported by the National Center for Research Resources of the National Institutes of Health under Award Number 1S20RR026802-1. The content is solely the responsibility of the authors and does not necessarily represent the official views of the NIH.

## Author contributions

K.G. and E.L.S. designed experiments. K.G., W.A.O., G.F., H.E.D., E.W., and X.Z. performed experiments. K.G., G.F., T.J.P., E.W., X.Z. and E.L.S. analyzed data. E.L.S. performed histopathologic review. K.G. and E.L.S. wrote the manuscript. All authors discussed results, reviewed, and revised the manuscript.

## Supplemental Tables

Table S1. DEGs in K, KN, KNF1F2, and N murine samples and TCGA Pan Cancer Atlas LUAD samples, related to Figures 1, S1, S2, 2, S3, 6, and S9.

Table S2. DMRs in K, KN, KNF1F2, and N murine samples, human methylation atlas cell types, organoids KG1A and 22E, and TCGA PanCancer LUAD samples, related to figures 1, S1, S2, 2, S3, S4, 5, S8, 6, and S9.

Table S3. Enrichr analysis on K, KN, KNF1F2, and N DMRs, differential H3K27ac in KN vs. KNF1F2, and DEGs in KN vs. N, related to figures 1, 2, 4, S6, 6, and S9.

Table S4. GSEA cell type (C8) and hallmark analysis on KNF1F2-specific and KG1A DMRs, related to figures 2, S4, 5, and S8.

Table S5. DEGs in KG1A and 22E organoid samples related to Figures 5 and S8.

## EXPERIMENTAL MODEL AND SUBJECT DETAILS

### Animal studies

Mice harboring *Kras^FSF-G12D^* (Young et al., 2011), *Rosa-FSF-Cre^ERT2^* (Schonhuber et al., 2014), *Nkx2-1^flox^*(Kusakabe et al., 2006), *Foxa1^flox^* (Gao et al., 2008), *Foxa2^flox^ (*Sund et al., 2000), *R26-CAG-LSL-Sun1-sfGFP-myc* (Mo et al., 2015), and *p53^frt^* (Lee et al., 2012), have been previously described. All animals were maintained on a mixed 129/B6 background. All experimental mice were between 2 and 6 months of age at intubation. Mice of both sexes were used throughout each study. Animal studies were approved by the IACUC of the University of Utah, conducted in compliance with the Animal Welfare Act Regulations and other federal statutes relating to animals and experiments involving animals, and adhered to the principles set forth in the Guide for the Care and Use of Laboratory Animals, National Research Council (PHS assurance registration number A-3031-01).

### Primary 3D organoid cultures

All primary murine organoid cultures (see key resources table) were established within Matrigel (Corning or Preclinical Research Shared Resource core facility) submerged in recombinant organoid medium for approximately two weeks (Advanced DMEM/F-12 supplemented with 1X B27 (Gibco), 1X N2 (Gibco), 1.25mM nAcetylcysteine (Sigma), 10mM Nicotinamide (Sigma), 10nM Gastrin (Sigma), 100ng/ml EGF (Peprotech), 100ng/ml R-spondin1 (Peprotech), 100ng/ml Noggin (Peprotech), and 100ng/ml FGF10 (Peprotech). After organoids were established, cultures were switched to 50% L-WRN conditioned media (Miyoshi and Stappenbeck, 2013). Organoid lines were tested periodically for mycoplasma contamination. To maintain organoid cultures mycoplasma free, all culture media were supplemented with 2.5 ug/ml Plasmocin.

## METHOD DETAILS

### Tumor initiation and tamoxifen administration in vivo

Autochthonous lung tumors were initiated by administering viruses via intratracheal intubation. Adenoviral mSPC-FlpO was used to initiate all tumors in this publication. Adenoviruses were obtained from University of Iowa Viral Vector Core.

Tumor-specific activation of Cre^ERT2^ nuclear activity was achieved by intraperitoneal injection of tamoxifen (Sigma) dissolved in corn oil at a dose of 120mg/kg. Mice received 4 injections over the course of 5 days. One day following injections, mice were given pellets supplemented with 500mg/kg (Envigo) tamoxifen for 7 days.

### Histology and immunohistochemistry

All tissues were fixed in 10% formalin overnight and when necessary, lungs were perfused with formalin via the trachea. Organoids were first fixed in 10% formalin overnight and then mounted in HistoGel (Thermo Fisher Scientific). Mounted organoids and tissues were transferred to 70% ethanol, embedded in paraffin, and four-micrometer sections were cut. Immunohistochemistry (IHC) was performed manually on Sequenza slide staining racks (Thermo Fisher Scientific). Sections were treated with Bloxall (Vector Labs) followed by Horse serum 536 (Vector Labs) or Rodent Block M (Biocare Medical), primary antibody, and HRP-polymer-conjugated secondary antibody (anti-Rabbit, Goat and Rat from Vector Labs; anti-Mouse from Biocare). The slides were developed with Impact DAB (Vector Labs) and counterstained with hematoxylin. Slides were stained with antibodies to NKX2-1 (1:2000, Abcam EP1584Y), GFP (1:200 CST 2956S) FoxA1 (1:4000, Abcam 10881-14), FoxA2 (1:1200, Abcam 4466), and HNF4a (1:500, CST C11F12). Images were taken on a Nikon Eclipse Ni-U microscope with a DS-Ri2 camera and NIS-Elements software. Histological analyses were performed on hematoxylin and eosin-stained and IHC-stained slides using NIS-Elements software. All histopathologic analysis was performed by a board-certified anatomic pathologist (E.L.S.).

### Establishing primary murine LUAD organoids

Five months after tumor initiation in KNF1F2 mice (3D organoid lines; Adeno-mSPC-FlpO), tumor bearing mice were euthanized and lungs were isolated. Individual macroscopic tumors were removed from lungs, minced under sterile conditions, and digested at 37°C for 30 min with continuous agitation in a solution of Advanced DMEM/F12 containing the following enzymes: Collagenase Type I (Thermo Fisher Scientific, 450U/ml), Dispase (Corning, 5U/ml), DNaseI (Sigma, 0.25mg/ml). Enzymatic reactions were stopped by addition of cold DMEM/F-12 with 10% FBS. The digested tissue was repeatedly passed through a 20-gauge syringe needle, sequentially dispersed through 100 mm, 70 mm, and 40 mm cell strainers, and treated with erythrocyte lysis buffer (eBioscience) to obtain a single cell suspension.

Organoid cultures were established by seeding 1x10^5^ tumor cells in 50ul of Matrigel (Corning) and plated in 24-well plates. Matrigel droplets were overlaid with recombinant organoid medium as previously described (Cell lines and primary cultures). Two weeks after organoid establishment, cultures were switched to 50% L-WRN conditioned media. Organoid cultures were screened via immunohistochemistry and qPCR, and lines that uniformly downregulated *Nkx2-1* but retained expression of *Foxa1* and *Foxa2* were selected for subsequent analysis. Of note, the 22E organoid line harbors one conditional allele of *Trp53* but maintains a well-differentiated gastric identity in the NKX2-1-negative state.

### In vitro 4-hydroxytamoxifen (4OHT) treatment

Cells were transiently treated with 2mM 4-OHT (Cayman Chemical Company, dissolved in 100% Ethanol) or vehicle for 72 (organoid culture) hr to activate Cre^ERT2^ nuclear activity and generate isogenic pairs.

### Generating a single cell suspension from organoid cultures

Matrigel droplets containing organoid cells were broken down via repeated pipetting in Cell Recovery Solution (Corning, 500 µl per Matrigel dome). Cell Recovery Solution containing organoids was transferred to sterile conical tubes and submerged in ice for 20–30 min before centrifugation at 4°C (300-500G). Cell Recovery Solution supernatant was removed and the cell pellet was washed in 1X PBS. Cells were then resuspended in pre-warmed TrypLE Express Enzyme (Thermo Fisher Scientific) and incubated for 10 min at 37°C. TrypLE reaction was quenched via dilution with cold Splitting Media (Advanced DMEM/F-12 [Gibco], 10 mM HEPES [Invitrogen], 1X Penicillin-Streptomycin-Glutamine [Invitrogen]). Cells were centrifuged and then resuspended in a pre-warmed DNase solution (L-WRN media supplemented to a final concentration of 200U/ml DNase [Worthington], 2.5 mM MgCl2, 500 mM CaCl2) and incubated for 10 min at 37°C. Cells were centrifuged and washed in PBS before use.

### DNA Methylation sequencing

#### In vitro Methyl-seq

DNA was collected from biological replicates of isogenic organoid cultures, KG1A and 22E. Four Matrigel domes per sample were collected two weeks after 4-OHT or ethanol treatment. Matrigel domes were resuspended directly into Cell Recovery Solution (500 µl per dome) and submerged in ice for 30 min before centrifugation at 4°C (300G). Cells were washed twice in cold PBS and cell pellets were frozen at -80°C.

DNA was purified from organoid cell pellets using the DNeasy Blood and Tissue kit according to the manufacturer’s instructions (Qiagen). Library preparation was performed using the NEBNext Enzymatic Methyl-seq Kit. Lambda phage DNA was added as a spike-in control for measuring conversion efficiency. Sequencing was performed using the Illumina NovaSeq 6000 (150 x 150 bp paired-end sequencing, 100 million reads per sample).

#### In vivo Methyl-seq

14 weeks after tumor initiation, K, N, KN and KNF1F2 mice were euthanized and the rib-cage was dissected to reveal the trachea and heart. Lungs were perfused with cold PBS, removed, and snap frozen in liquid nitrogen. Flash-frozen lungs were minced on ice in 1 ml of ice-cold Tween with salts and Tris (TST) buffer^55^ (146 mM NaCl, 10 mM Tris-HCL pH 7.5, 1 mM CaCl_2_, 21 mM MgCl_2_, 0.01% BSA, 0.006% Tween-20 in ultrapure water) for 5 – 8 min depending on tumor burden. Lysis was quenched with 3 ml of 1X salts and Tris (ST) buffer (146 mM NaCl, 10 mM Tris-HCL pH 7.5, 1 mM CaCl_2_, 21 mM MgCl_2_ in ultrapure water) and solution was filtered through a 40 µm cell strainer. Sample was transferred to a 15 ml conical and centrifuged for 5 min at 4°C (300G). Nuclei pellet was carefully resuspended in 1 ml of FACS buffer (1X PBS, 1% BSA, 1% serum, 2 mM EDTA) with protease inhibitors and filtered through a 35 µm cell strainer. Samples were stained with DAPI (Sigma) and evaluated on a fluorescence microscope to assess for GFP positivity and nuclear integrity. Nuclei were sorted on the BD FACSAria with the 100 μm nozzle to obtain a GFP-positive, DAPI-positive population. Samples were sorted into 1 ml of cold PBS with 10% serum. After sorting, nuclei were centrifuged for 10 min at 4°C (300G) and pellets were frozen at -80°C. DNA was purified and sequenced as previously described (In vitro Methyl-seq).

#### Methyl-seq data processing and analysis

Fastq reads were aligned to the mouse genome (build mm10) with Novocraft novoalign (version 4.03.01) in bisulfite mode (-b 4) with the following options: set penalty for unconverted CHG or CHH cytosines (option -u) to 12, hard clip 3’ bases to quality 20 (option -H), adapter trimming (option -a), and performance tuning set to ‘NOVOSEQ’. Alignments to non-standard chromosomes were ignored. Extreme coverage regions were identified using depth-normalized (Reads Per Million) mean coverage of replicates and MACS2 bdgpeakcall with an absolute depth cutoff of 2 RPM (mean depth was < 0.1), length 200 bp, and gap 500 bp. Alignments overlapping these exclusion intervals were excluded from further analysis. Duplicate alignments were removed using samtools (version 1.16) fixmate and markdup (mode -s).

Alignments were processed with USeq NovoalignBisulfiteParser (version 9.3.0) to collect converted and non-converted C counts with minimum mapping quality score (-q) of 13 and minimum base quality score (options -b and -c) of 20. Counts in CpG context were extracted with USeq ParsePointDataContexts using the context “..CG.”. Fraction methylation tracks were generated using USeq BisStat requiring a minimum coverage of 4 reads. Measured lambda phage DNA conversion efficiency was typically > 99.7%. BisMark compatible coverage files were generated from the Converted and NonConverted .useq files using the useq2bismark_methylation_extractor application (https://github.com/tjparnell/HCI-Scripts/blob/master/Methylation/useq2bismark_methylation_extractor.pl).

DMRs were identified following the recommended pipeline detailed in the bsseq^31^ package. CpG counts were smoothed using BSmooth and default parameters of 70 CpGs and sliding window of 1000 bp. CpGs were subsequently filtered for a minimum coverage of at least 5 reads. For each pairwise comparison, t-statistic values (differences in means) were generated at each CpG. Regions with differential CpGs were identified with dmrFinder using reciprocal quantile cutoffs of 2.5% (rather than absolute thresholds), a minimum number of 5 CpGs, a maximum gap of 200 bp, and an absolute minimum mean fraction methylation differential of 15%. DMRs were annotated to the nearest gene using the ChIPseeker package (citation) and custom annotation consisting of protein-coding, Gencode-basic genes from Ensembl annotation release 102.

For comparative analysis between multiple pair-wise comparisons, DMRs were intersected to generate a master list of all possible DMR intervals, and these were re-scored for mean CpG fraction methylation values (non-smoothened) with BioToolBox get_datasets (https://github.com/tjparnell/biotoolbox). To identify FoxA1/2 dependent or independent regions from the merged intervals, differential values were re-calculated from replicate mean values and filtered for the minimum change of 15% in the appropriate direction. Relative difference methylation levels were calculated in heat maps by subtracting the mean methylation value across all replicates. Heat maps were generated with pHeatmap package (citation). Motif analysis was performed on DMRs for both known and novel motifs using the HOMER package^56^. GSEA-Preranked was run on the differentially methylated gene lists generated from bsseq and ranked by their log2 fold change in methylation. DMGs were run against the following MSigDB gene sets: c2, c5, c6, c8, and Hallmarks. Gene sets smaller than 15 and larger than 500 were excluded from analysis.

### RNA sequencing

#### In vitro RNA-seq

RNA was collected from biological replicates of isogenic organoid cultures, KG1A and 22E. Samples were collected two weeks after 4-OHT or ethanol treatment. Three Matrigel domes were collected per sample directly into Trizol, then stored at -80°C until purification.

RNA was isolated via Trizol-chloroform extraction followed by column-based purification. The aqueous phase was brought to a final concentration of 50% ethanol, and RNA was purified using the PureLink RNA Mini kit according to the manufacturer’s instructions (ThermoFisher Scientific). Library preparation was performed using the NEBNext Ultra II Directional RNA Library Prep with poly(A) mRNA isolation. Sequencing was performed using the Illumina NovaSeq 6000 (150 x 150 bp paired-end sequencing, 25 million reads per sample).

#### In vivo RNA-seq

14 weeks after tumor initiation, K, N, KN, and KNF1F2 mice were euthanized and the rib-cage was dissected to reveal the trachea and heart. Cardiac perfusion of the pulmonary vasculature was performed using PBS until the lungs turned pale. Lungs were then removed, digested, and filtered as previously described (Establishing primary murine LUAD organoids) to obtain a single cell suspension. Samples were resuspended in FACS buffer with DAPI. Cells were sorted on the BD FACSAria with the 85 μm nozzle to obtain a GFP-positive, DAPI-negative population. Samples were sorted into 1 ml of cold PBS with 10% serum. After sorting, cells were centrifuged for 10 min at 4°C (300G) and resuspended in 1 ml of Trizol. RNA was isolated via Trizol-chloroform extraction and the PureLink RNA Mini kit as previously described (In vitro RNA-seq). Library preparation was performed using the NEBNext Ultra II Directional RNA Library Prep with rRNA Depletion Kit for mouse. Sequencing was performed using the Illumina NovaSeq 6000 (150 x 150 bp paired-end sequencing, 25 million reads per sample).

#### RNA-seq data processing and analysis

The mouse GRCm38 genome and gene feature files were downloaded from Ensembl release 102 and a reference database was created using STAR version 2.7.6a (Dobin et al., 2013). Optical duplicates were removed from NovaSeq runs via Clumpify v38.34 (Bushnell, 2021). Reads were trimmed of adapters and aligned to the reference database using STAR in two-pass mode to output a BAM file sorted by coordinates. Mapped reads were assigned to annotated genes using featureCounts version 1.6.3 (Liao et al., 2019). Raw counts were filtered to remove features with zero counts and features with five or fewer reads in every sample. DEGs were identified using the hciR package (https://github.com/HuntsmanCancerInstitute/hciR) with a 5% false discovery rate and DESeq2 version 1.34.0 (Love et al., 2014). GSEA-Preranked was run with the differential gene list generated from DESeq2 and the following MSigDB gene sets: c2, c5, c6, c8, and Hallmarks. Gene sets smaller than 15 and larger than 500 were excluded from analysis. MEK activation signature^57^ was downloaded from Dry et al. supplemental materials. ChEA3 was performed on DEGs identified from DESeq2 (log_2_FC > 1; padj < 0.05).

#### TCGA analysis of human lung adenocarcinoma

We first filtered all TCGA Pan Cancer Atlas lung adenocarcinoma samples by the presence of *KRAS* mutations and availability of expression data (n=154/562 LUAD samples as of August 2023). The 154 KRAS-mutant samples were further filtered on *NKX2-1* and *HNF4A* expression. We first selected patients with above average *NKX2-1* expression (z-score>0.5; n=85/154) and low *HNF4A* expression (z-score<-1, n=41/85) to comprise the pulmonary K tumor cohort. We then selected patients with very low *NKX2-1* expression (z-score<-2; n=25/154) to capture tumor samples that have completely downregulated *NKX2-1*. We then filtered this group by high HNF4a expression (z-score>3, n=15/25) to comprise the gastric KN cohort. GSEA-Preranked was ran on human K and KN groups using the gene expression log2 ratio values generated by TCGA.

### Chromatin immunoprecipitation sequencing

#### In vitro organoid ChIP-seq

For all organoid ChIP-seq experiments, two 24-well plates (approximately 4 – 8 million cells) were collected in Cell Recovery Solution (500 µl per dome). Organoids were submerged in ice for 30 min before centrifugation at 4°C (300G). Cells were washed three times in cold PBS. On the second wash, PBS was supplemented with DNase solution (containing a final concentration of 200U/ml DNase [Worthington], 2.5 mM MgCl2, 500 mM CaCl2). After the third wash, organoids were resuspended in 5 ml of 2 mM DSG buffer (1X PBS, 1 mM MgCl_2_) and rotated at room temperature for 35 min. Formaldehyde was then added to a final concentration of 1% and cells were crosslinked for 10 min. The cross-linking reaction was stopped with the addition of glycine to a final concentration of 125 mM. Cells were washed with cold PBS, then frozen at -80. Cell pellets were thawed on ice for 5 min then lysed in 1 ml of Farnham lysis buffer (5 mM PIPES pH 8.0, 85 mM KCl, 0.5% NP40) for all TF experiments or 400 ul of Chromatrap Hypotonic buffer (catalog #100007) for all histone experiments. Samples were centrifuged at 4°C (1000G) then resuspended in 1ml of RIPA lysis buffer (1X PBS, 1% NP40, 0.5% sodium deoxycholate, 0.1% sodium dodecyl sulfate) or 100 µl of Chromatrap Lysis buffer (catalog #100005) for TF and histone ChIPs respectively. All lysis buffers were supplemented with protease inhibitors. Chromatin extract was sonicated with QSonica Q800R (pulse: 30 s on/30 s off; sonication time: 20 min; amplitude: 70%). After sonication, chromatin extract of histone samples was brought up to a volume of 1 ml with RIPA lysis buffer. Chromatin from all samples was immunoprecipitated with antibodies premixed with Protein A Dynabeads: FoxA1 (Abcam, ab170933, rabbit monoclonal, 5 µg/ChIP).

Library preparation was performed using the ChIP-seq with NEBNext DNA Ultra II library prep kit using Unique Molecular Indexes (UMIs). Sequencing was performed using the Illumina NovaSeq 6000 (150 x 150 bp paired-end sequencing, 25 million reads per sample). For ChIP-qPCR assays, primers were designed targeting FoxA1 binding sites identified from ChIP-seq data on Integrative Genomics Viewer (IGV; https://software.broadinstitute.org/software/igv/). All the primers are listed in Table S6.

#### In vivo nuclei ChIP-seq

14 weeks after tumor initiation, KN and KNF1F2 tumor-bearing mice were euthanized and the rib-cage was dissected to reveal the trachea and heart. Lungs were perfused with cold PBS, removed, and snap frozen in liquid nitrogen. Flash-frozen lungs were minced on ice in 2 ml of ice-cold PBS for 3 – 5 min. Minced lungs were dounced 10 times with a large 7 ml homogenizer (Wheaton, item #23ND78) and centrifuged for 5 min at 4°C (300G). Tissue was crosslinked in 2 mM DSG buffer and 1% Formaldehyde as previously described (In vitro organoid ChIP-seq). After fixation, lung samples were resuspended in 2 – 3 ml of TST buffer with protease inhibitors for 5 min. During this time, samples were dounced an additional 5 times in a small 2 ml homogenizer (Wheaton, item #23ND70) to extract nuclei. Lysis was quenched with 5 ml of 1X ST buffer plus protease inhibitors. Nuclei were washed in cold PBS and resuspended in 5 – 10 ml cold PBS supplemented with DAPI and protease inhibitors. Nuclei suspension was then sequentially filtered through a 70 μm and 35 μm cell strainer. Before sorting, nuclei were evaluated on a fluorescence microscope to assess for GFP positivity and nuclear integrity. Nuclei were sorted using BD FACSAria with the 85 μm nozzle for GFP-positive, DAPI-positive nuclei. Samples were sorted into 1 ml of cold PBS with 1% BSA and 10X protease inhibitor. Approximately 3 million nuclei were sorted for histone ChIP-seq and 10 million for TF ChIP-seq experiments.

After sorting, nuclei were pelleted for 10 min at 4°C (500G). Samples were then resuspended in Chromatrap Hypotonic and Lysis buffers as previously described (In vitro organoid ChIP-seq) and sonicated with QSonica Q800R (pulse: 30 s on/30 s off; sonication time: 20 min; amplitude: 70%). Chromatin was immunoprecipitated with antibodies premixed with Protein A Dynabeads: FoxA1 (Abcam, ab170933, rabbit monoclonal, 5 µg/ChIP), TET3 (Sigma, ABE290, rabbit polyclonal, 5 µg/ChIP) and H3K27ac (Active Motif, 39133, rabbit polyclonal, 5 µg/ChIP). ChIP-seq libraries were prepped and sequenced as previously described (In vitro organoid ChIP-seq). For ChIP-qPCR assays, primers were designed targeting FoxA1, TET3, and H3K27ac binding sites identified from ChIP-seq data on IGV. All the primers are listed in Table S6.

#### ChIP-seq data processing and analysis

Fastq alignments were pre-processed with the merge_umi_fastq application from the UMIScripts package (https://github.com/HuntsmanCancerInstitute/UMIScripts) to associated the UMI sequence, provided as a third Fastq file, into the read comment. Reads were aligned using Bowtie2^58^ to the standard chromosomes of the mouse genome (version mm10). Duplicate alignments based on the UMI code were removed using the bam_umi_dedup application (UMIScripts) allowing for 1 mismatch. Peaks were called using MACS2^59^ with a significance of q-value < 0.05. Coverage tracks were generated with MACS2 as Reads Per Million. Input libraries were obtained from all tumor and organoid samples and were used as controls for each ChIP-seq experiment. All ChIP-seq experiments were performed in biological duplicates. Genomic annotation of binding sites was performed using the HOMER package^56^. Overlaps between peaks were determined using Bedtools^60^ with a 1-bp minimum overlap, and percentages were calculated by dividing the number of overlapping peaks by the number of peaks in the smaller set (i.e., the percentage of maximal possible overlap). Motif finding was performed on 100-bp regions surrounding the summit of identified peaks. Both known and novel motifs were identified using the Homer package^56^. Differential ChIP-seq peaks were identified using the Diffbind package (https://bioconductor.org/packages/release/bioc/vignettes/DiffBind/inst/doc/DiffBind.pdf) with a q-value cutoff < 0.05. Motif finding for differential ChIP-seq peaks was performed by extending the differential region to a length of 1000 bps. HOMER motif analysis was then performed on the extended differential region to identify known and novel motifs. Heat maps and plots were generated with custom R scripts using pHeatmap (https://cran.r-project.org/ package=pheatmap) and ggplot2 (https://ggplot2.tidyverse.org).

#### Hi-C chromatin immunoprecipitation (HiChIP)

Tumor processing, fixation, and sorting was performed as previously described (“In vivo nuclei ChIP-seq”). HiChIP was performed as described (https://www.nature.com/articles/nmeth.3999) with minor modifications as described (https://www.nature.com/articles/s41467-021-27055-4). Cross-linked chromatin was digested with the MboI restriction enzyme followed by end-repair with dNTPs including biotin labeled dATP, ligation using T4 DNA ligase, and sonication to obtain 1kb chromatin fragments using Qsonica (Q800). To enrich for chromatin interactions occurring at active regulatory elements, an anti-H3K27ac antibody (Abcam, ab4729, rabbit polyclonal, 7.5ug/HiChIP) was used for DNA fragment capture. Streptavidin magnetic beads were used to pull down ligated DNA fragments and HiChIP libraries were prepared using Illumina Tagment DNA Enzyme and Buffer Kit. Sequencing was performed with Illumina NextSeq.

The paired-end HiChIP sequencing reads were aligned to the mouse genome mm10 with the HiC-Pro pipeline (https://genomebiology.biomedcentral.com/articles/10.1186/s13059-015-0831-x). HiC-DC+ was used to call chromatin loops by binning the genome into 5kb bins (https://www.nature.com/articles/s41467-021-23749-x#citeas). Chromatin loops were removed if the qvalue < 0.05 or anchors overlapped with ENCODE blacklist regions (https://www.nature.com/articles/s41598-019-45839-z). BEDTools pairToBed was used to identify enhancer to promoter interactions (https://academic.oup.com/bioinformatics/article/26/6/841/244688). HiChIP data was visualized at the resolution of 5kb using the R package gTrack (https://github.com/mskilab-org/gTrack).

## QUANTIFICATION AND STATISTICAL ANALYSIS

All graphing and statistical analysis was performed with PRISM software, with all graphs showing mean and standard deviation. The statistical details can be found in the corresponding figure legend. All NGS statistical analysis was performed according to published pipeline protocols cited, with a statistical significance cutoff of padj<0.05.

## RESOURCE AVAILABILITY

### Lead contact

Further information and requests for resources and reagents should be directed to and will be fulfilled by the lead contact, Eric Snyder

### Materials availability

Novel murine organoid lines are available upon request.

### Data and code availability

Bulk RNA-seq, Methyl-seq, ChIP-seq, and Hi-ChIP data have been deposited at Gene Expression Omnibus GSEXXXXXX.

---

**Supplement 1, related to Figure 1:**
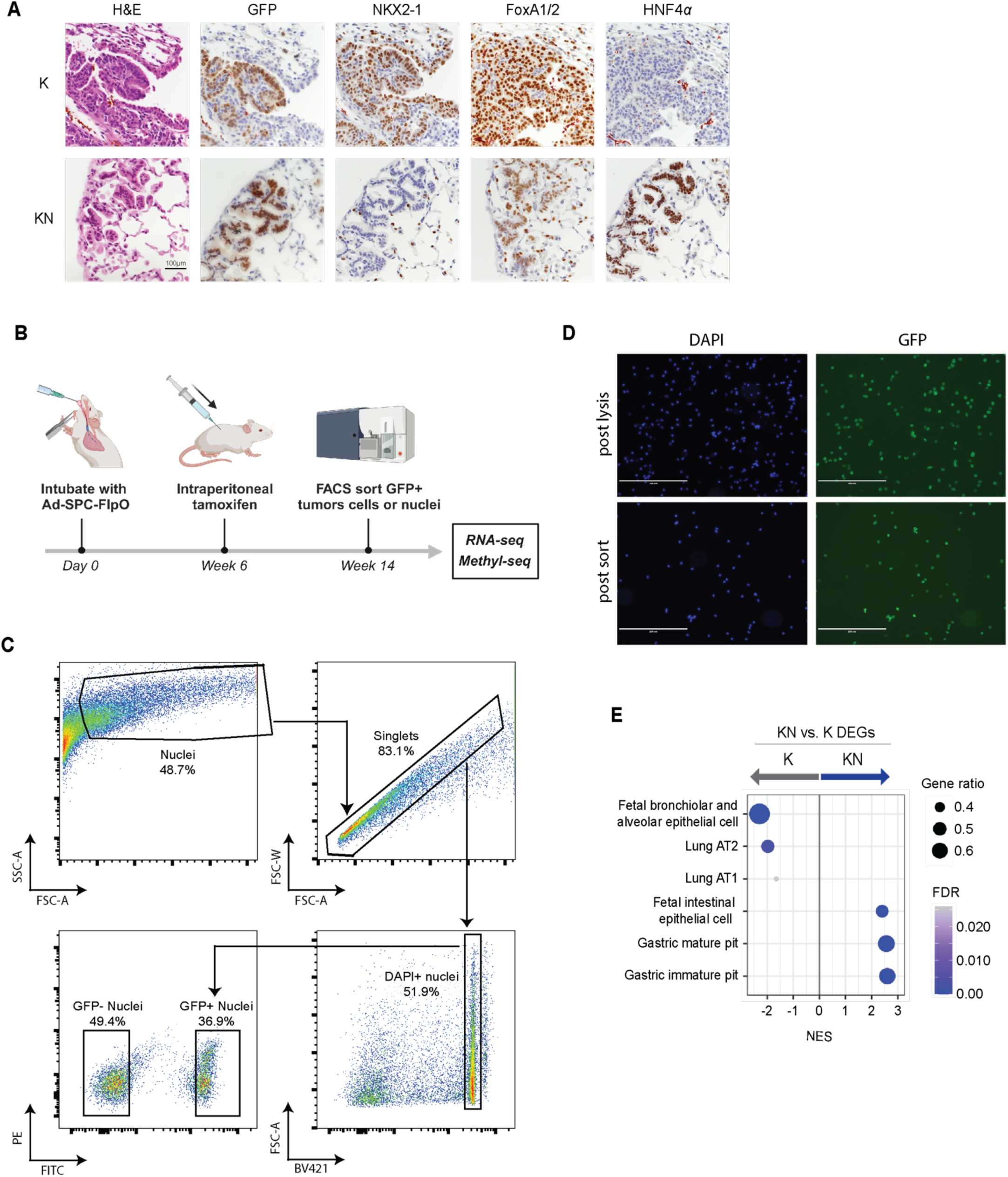
A) Representative images of K and KN tumors. IHC of GFP, NKX2-1, HNF4a, and hematoxylin and eosin (H&E) shown. B) Schematic of experimental timepoints for collection of tumor cells and nuclei. C) Representative FACS sorting strategy to isolate GFP+ DAPI+ nuclei from bulk lung. D) Fluorescent images of DAPI stained nuclei and Sun1-GFP expression from in vivo KRAS-driven LUAD tumors following nuclei extraction (post-lysis) and FACS sorting of GFP+ nuclei (post-sort). Magnification 200um. E) GSEA for cell type signatures on DEGs identified between K and KN tumors.

**Supplement 2, related to Figure 1:**
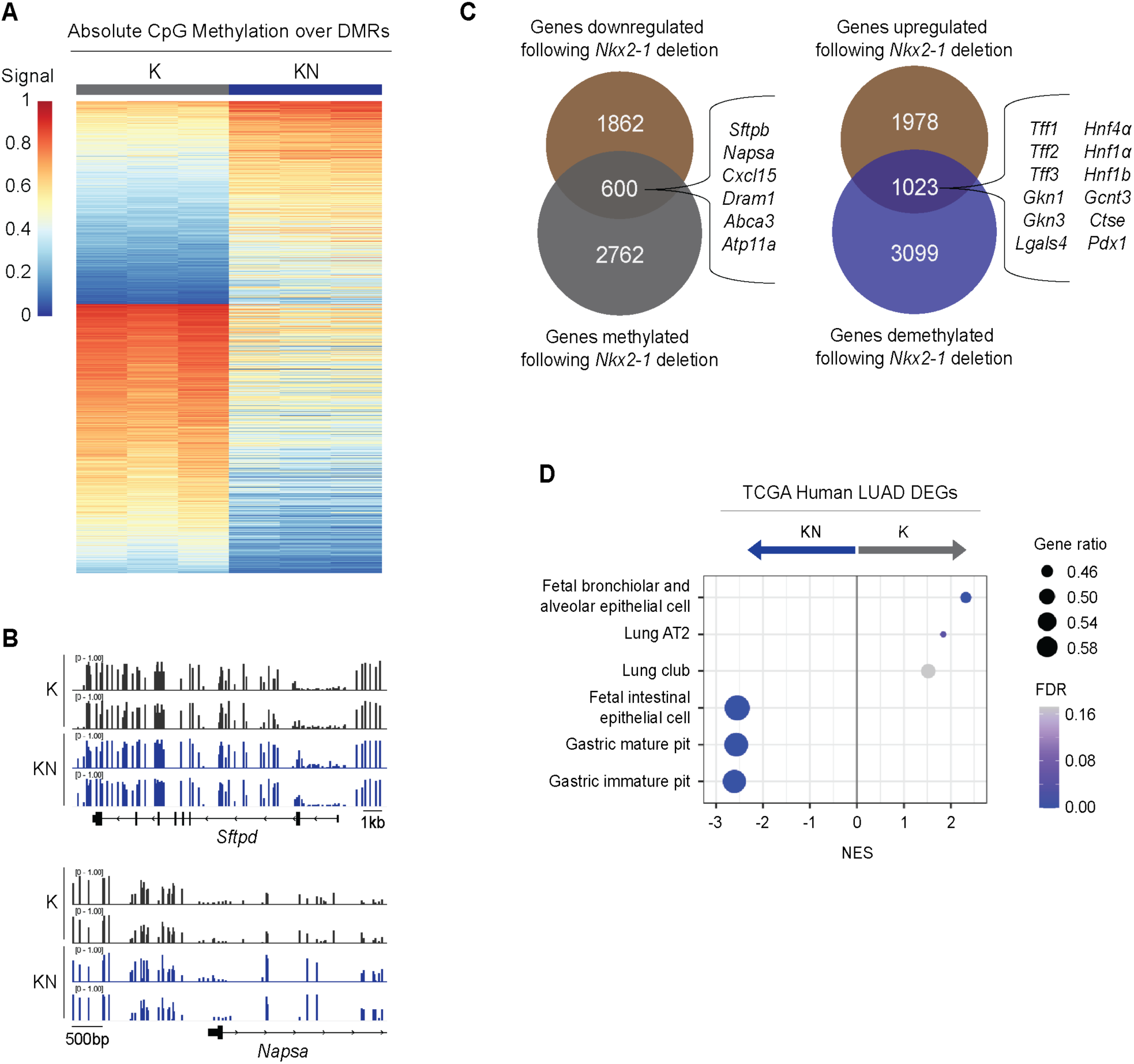
A) Heatmap of DMRs identified in K and KN tumors showing absolute difference in mean methylation at CpGs. DNA methylation scores calculated by subtracting the row mean from the average methylation level of each DMR per sample. B) Methylation tracks at AT2 genes, *Sftpd* and *Napsa*, that do not undergo methylation changes in K vs. KN tumors. C) Intersection of DEGs and DMGs in K and KN tumors. Shown are representative pulmonary (blue) or gastric (red) genes where gene expression correlates with methylation. D) GSEA for cell type signatures on DEGs identified from K and KN human LUAD patients (TCGA Pan Cancer Atlas).

**Supplement 3, related to Figure 2:**
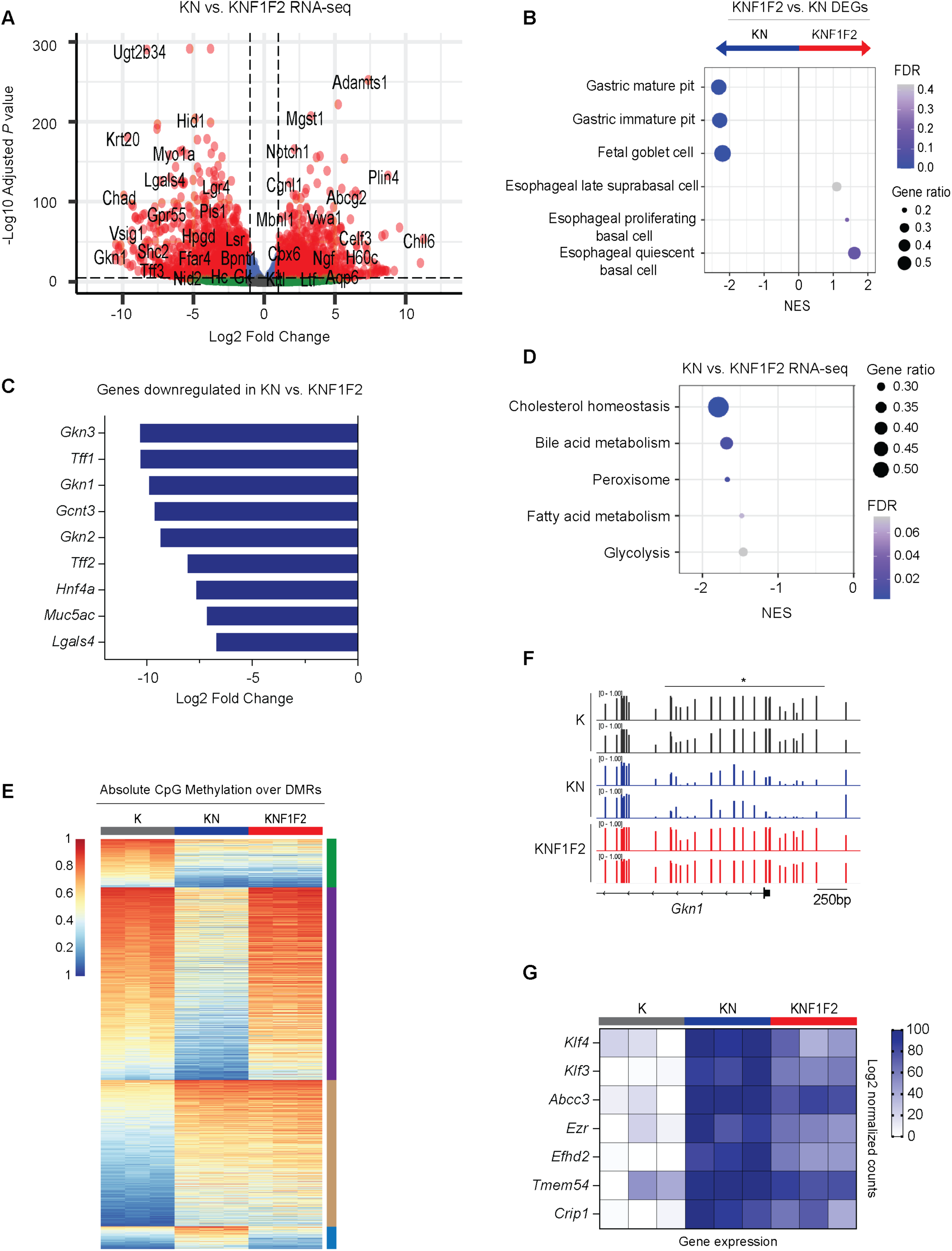
A) Volcano plot of DEGs identified between KN and KNF1F2 tumors. Plotted are the log2 fold change values against -log_10_ adjusted *p*-value. Genes indicated by red dots have a p-value < 0.05 and log2 fold change > 1. B) GSEA for cell type signatures on DEGs identified between KN and KNF1F2 tumors. C) Log2 fold change values of GI genes downregulated with *FoxA1/2* deletion in KN tumors. D) GSEA for hallmark signatures on DEGs identified between KN and KNF1F2 tumors. E) Heatmap showing absolute difference in mean methylation of CpGs at DMRs identified in K, KN, and KNF1F2 tumors. DMRs categorized by FoxA1/2-dependency and methylation status (e.g., methylated vs. demethylated). F) Methylation tracks at gastric pit cell gene, *Gkn1*, in K, KN, and KNF1F2 tumors. Lines with asterisk indicate significant DMRs. G) Heatmap showing log2 normalized counts of genes comprising the gastric pit cell signature in K, KN, and KNF1F2 samples. Genes shown contained both FoxA1/2-dependent and independent hypoDMRs.

**Supplement 4, related to Figure 2:**
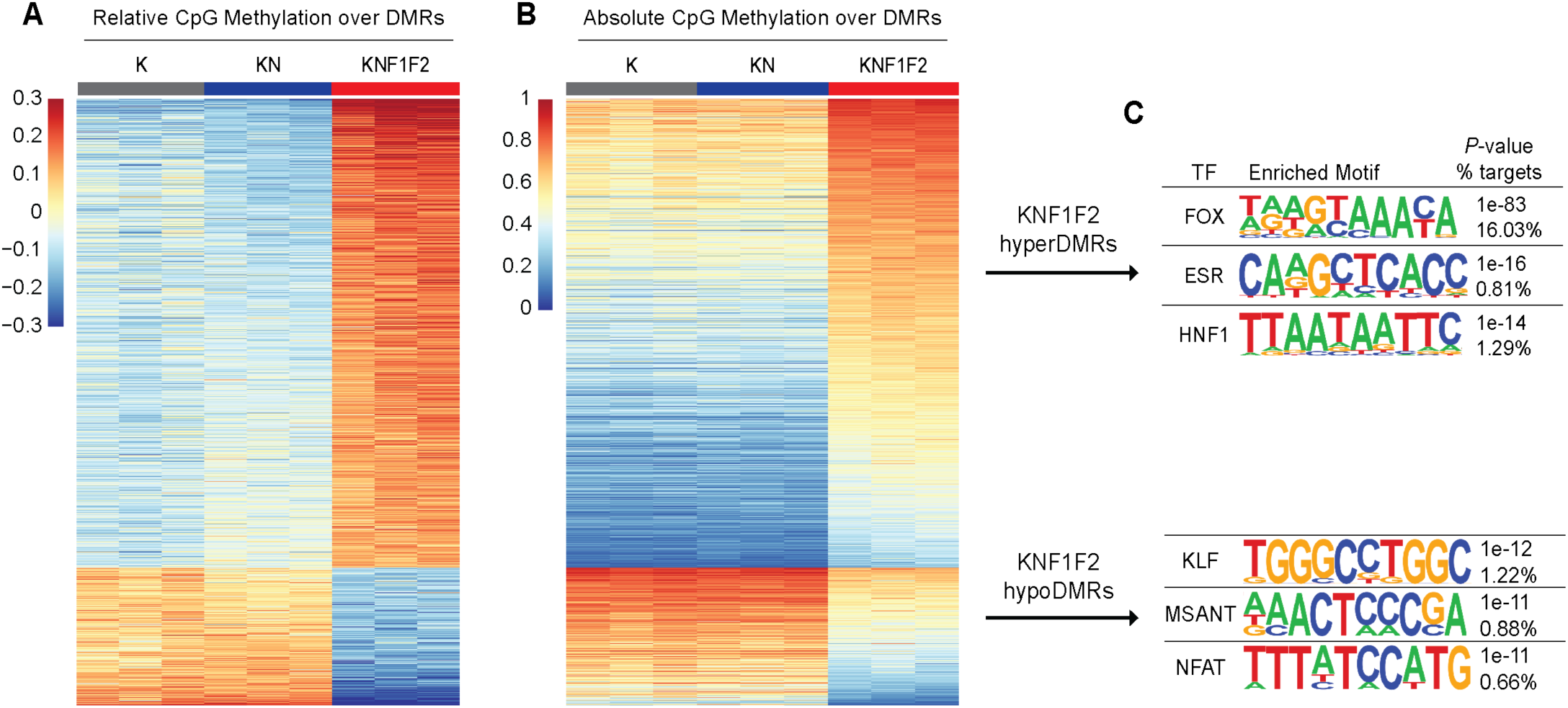
A) Heatmap showing relative difference in mean methylation of CpGs at DMRs specific to KNF1F2 tumors. Shown are K, KN, and KNF1F2 samples. B) Heatmap showing absolute difference in mean methylation of CpGs at DMRs specific to KNF1F2 tumors. Shown are K, KN, and KNF1F2 samples. C) TF motifs enriched in KNF1F2-specific hyper- vs. hypoDMRs.

**Supplement 5, related to Figure 3:**
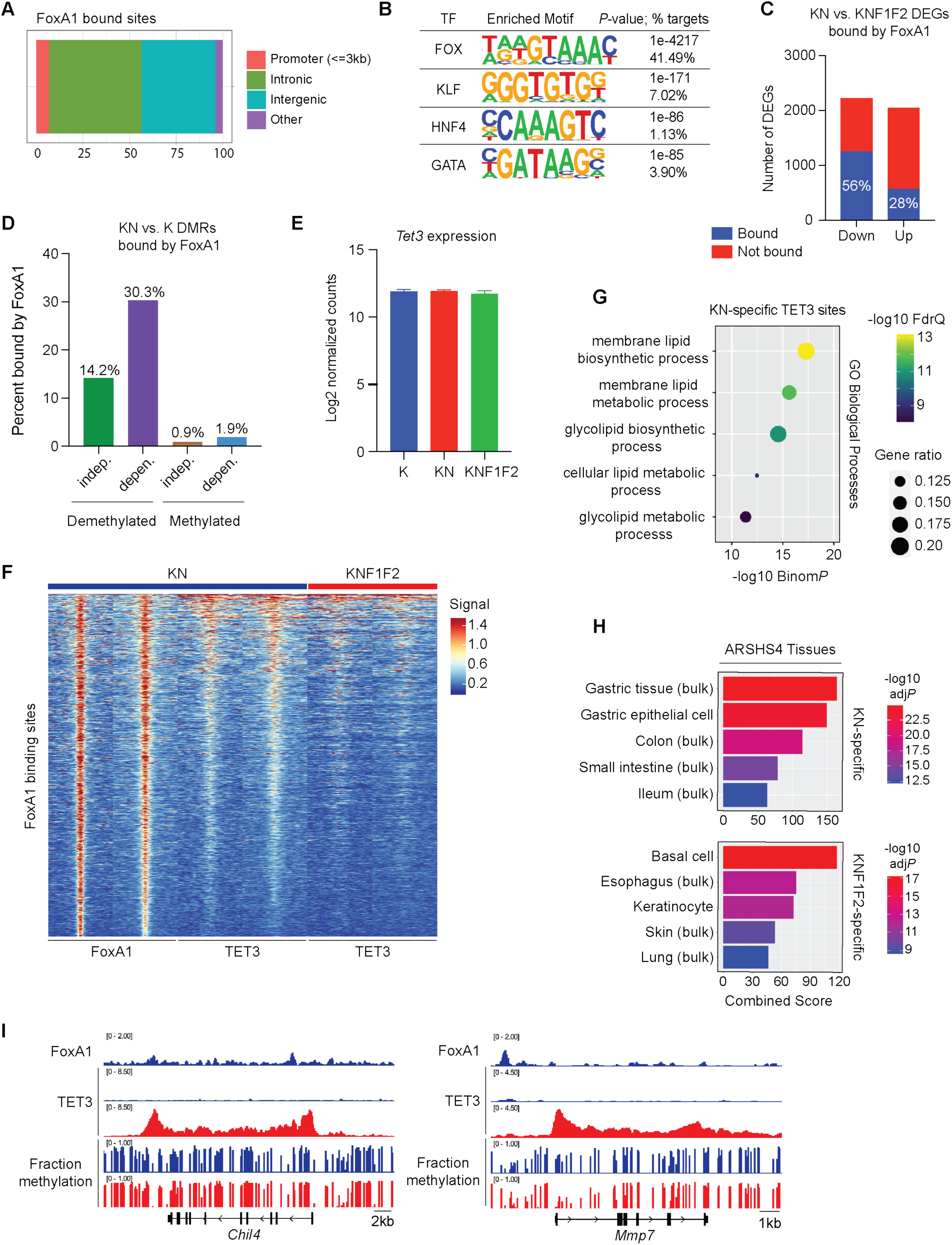
A) Feature distribution of FoxA1 bound sites in KN tumors. B) TF motifs enriched at FoxA1 bound sites in KN tumors. C) Number of DEGs identified in KN and KNF1F2 tumors that contain a FoxA1 binding site. FoxA1 sites overlaps significantly more with downregulated genes (Fisher’s exact test; p-value < 10^-4^). D) Intersection of FoxA1 ChIP peaks with DMRs defined in Figure 2B. FoxA1 binding sites are significantly associated with DMRs demethylated in a FoxA1/2-dependent manner (Fisher’s exact test; p-value < 10^-4^). E) Log2 normalized counts of *Tet3* expression in K, KN, and KNF1F2 tumors. F) Heatmap of FoxA1 and TET3 occupancy in KN and KNF1F2 tumors at 17,822 FoxA1 bound sites. G) GO biological processes enriched at KN-specific TET3 bound sites. H) Cell type analysis with ARCHS4 tissue database using Enrichr on KN and KNF1F2 differential TET3 bound sites. I) FoxA1 and TET3 ChIP peaks and methylation tracks at SCJ genes, *Chil4* and *Mmp7*, in KN (blue) and KNF1F2 (red) tumors.

**Supplement 6, related to Figure 4:**
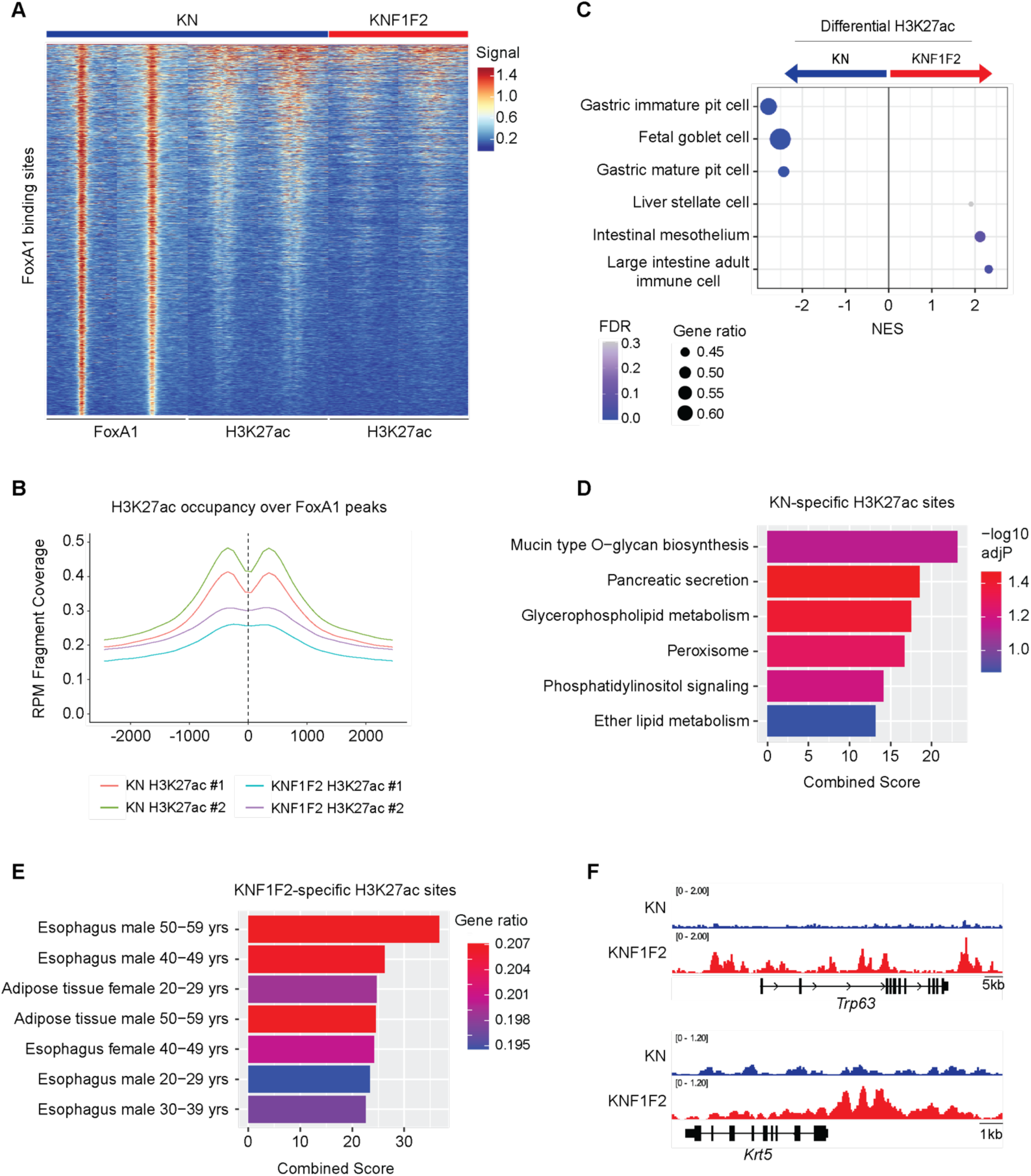
A) Heatmap of FoxA1 and H3K27ac occupancy in KN and KNF1F2 tumors at 17,822 FoxA1 bound sites. B) Mean occupancy profile of H3K27ac at FoxA1 bound sites in KN and KNF1F2 tumors. C) GSEA for cell type signatures on differentially bound H3K27ac sites identified between KN and KNF1F2 tumors. D) Pathway analysis with the KEGG database using Enrichr on KN-specific H3K27ac bound sites. E) Cell type analysis with the GTEx database using Enrichr on KNF1F2-specific H3K27ac bound sites. F) H3K27ac ChIP peaks at squamous genes, *Trp63* and *Krt5*, in KN (blue) and KNF1F2 (red) tumors.

**Supplement 7, related to Figure 4:**
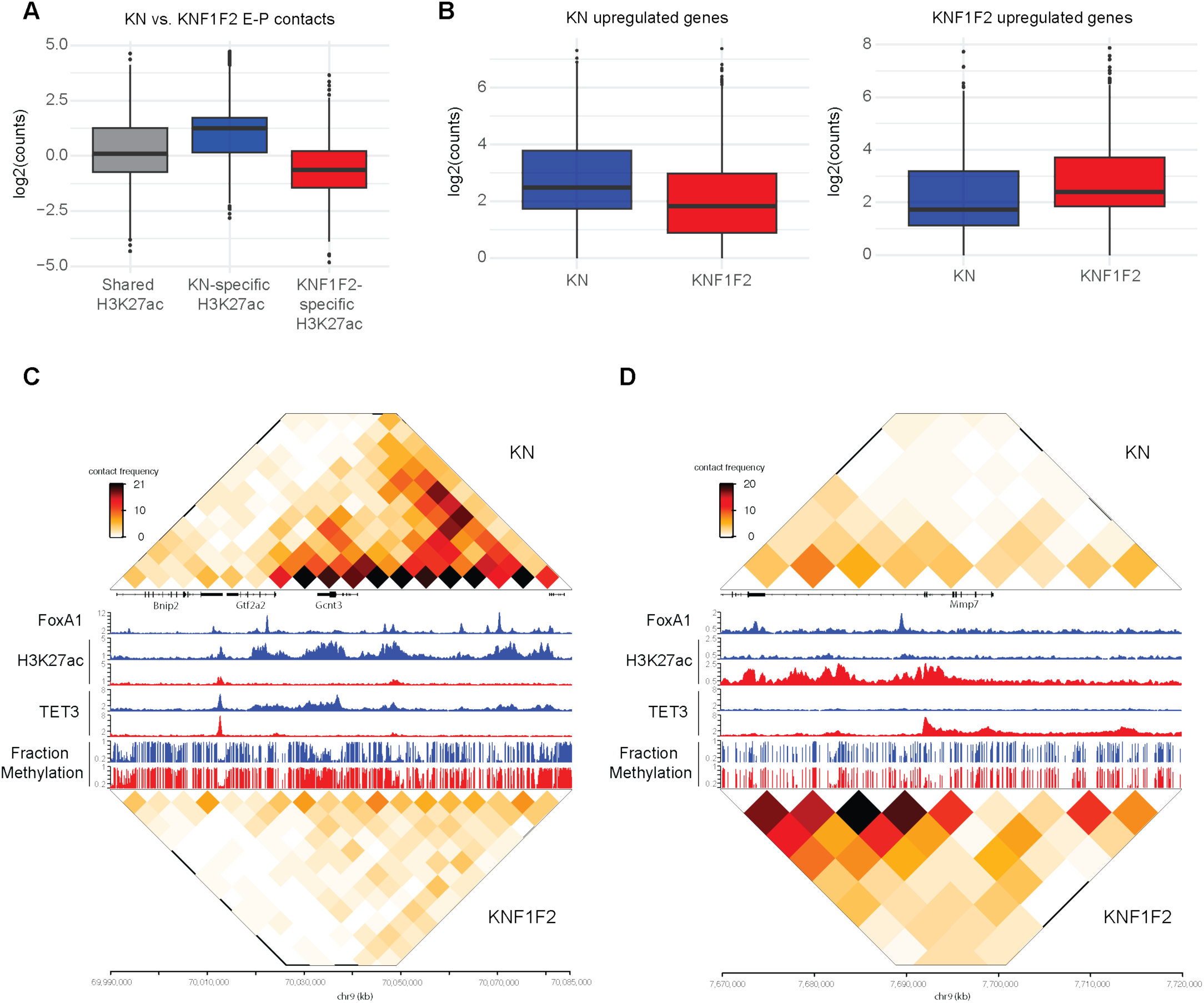
A) Log2 counts of enhancer-promoter contacts in KN vs. KNF1F2 tumors at shared, KN-specific, and KNF1F2-specific H3K27ac sites. B) Log2 counts of enhancer-promoter contacts in KN vs. KNF1F2 tumors at genes upregulated in KN and KNF1F2 tumors. C) Heatmap of H3K27ac contact frequency with FoxA1, TET3, and H3K27ac ChIP peaks and methylation tracks at pit cell marker, *Gcnt3*, in KN (blue) and KNF1F2 (red) tumors. D) Heatmap of H3K27ac contact frequency with FoxA1, TET3, and H3K27ac ChIP peaks and methylation tracks at SCJ marker, *Mmp7*, in KN (blue) and KNF1F2 (red) tumors.

**Supplement 8, related to Figure 5:**
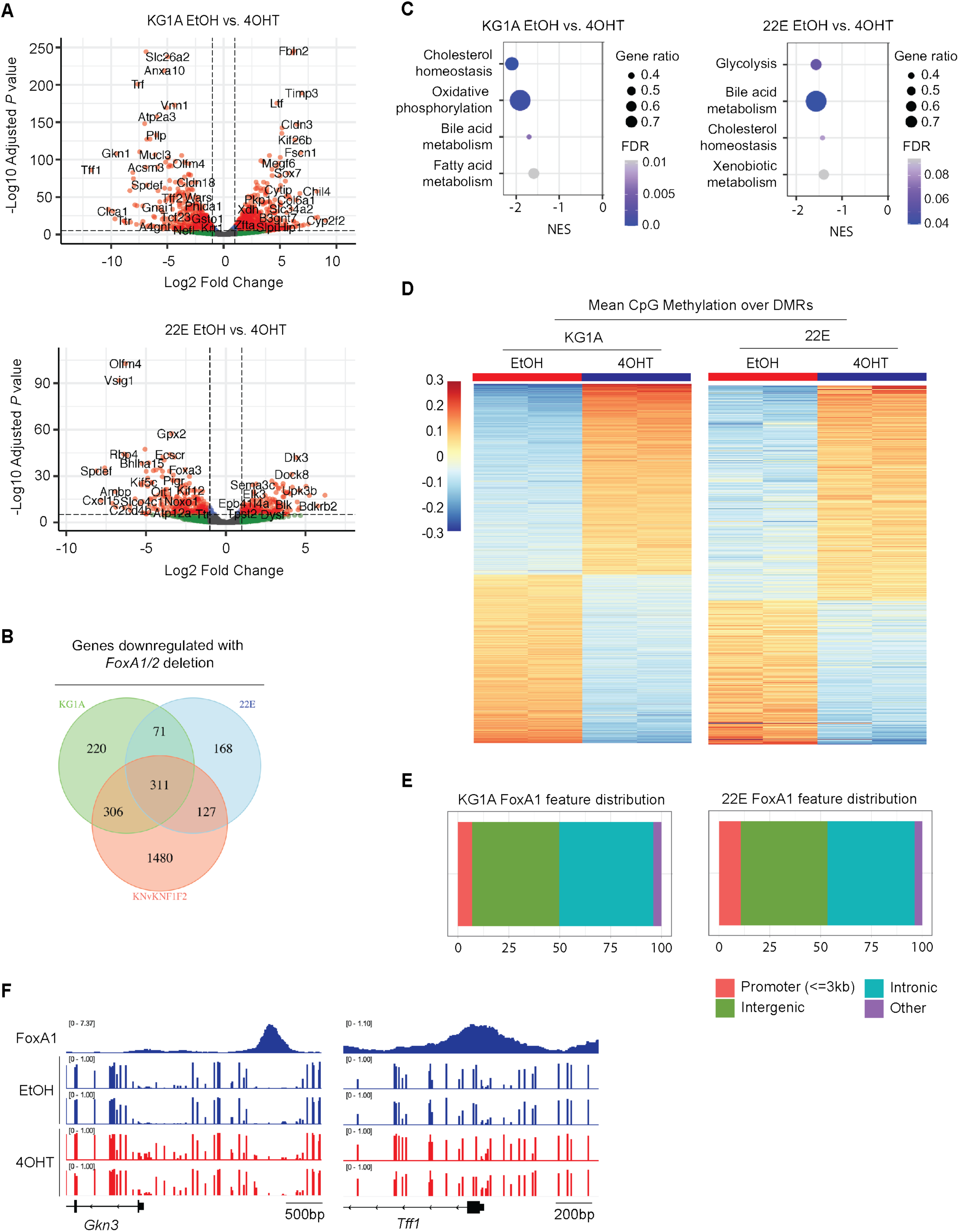
A) Volcano plot of DEGs identified in EtOH vs. 4OHT for KG1A and 22E organoids. Plotted are the log2 fold change values against -log_10_ adjusted *p*-value. Genes indicated by red dots have a p-value < 0.05 and log2 fold change > 1. B) Venn diagram showing overlap of genes downregulated with *Foxa1/2* deletion in KG1A and 22E organoids and KN tumors. C) GSEA for hallmark signatures on DEGs identified in EtOH vs. 4OHT for KG1A and 22E organoids. D) Heatmap of mean relative CpG methylation over DMRs identified in EtOH vs. 4OHT for KG1A and 22E organoids. Samples collected two weeks post treatment. E) Feature distribution of FoxA1 bound sites in KG1A and 22E organoids treated with EtOH. F) FoxA1 ChIP peaks and methylation tracks at gastric genes, *Gcnt3* and *Tff2*, in KG1A organoids treated with EtOH (blue) or 4-OHT (red).

**Supplement 9, related to Figure 6:**
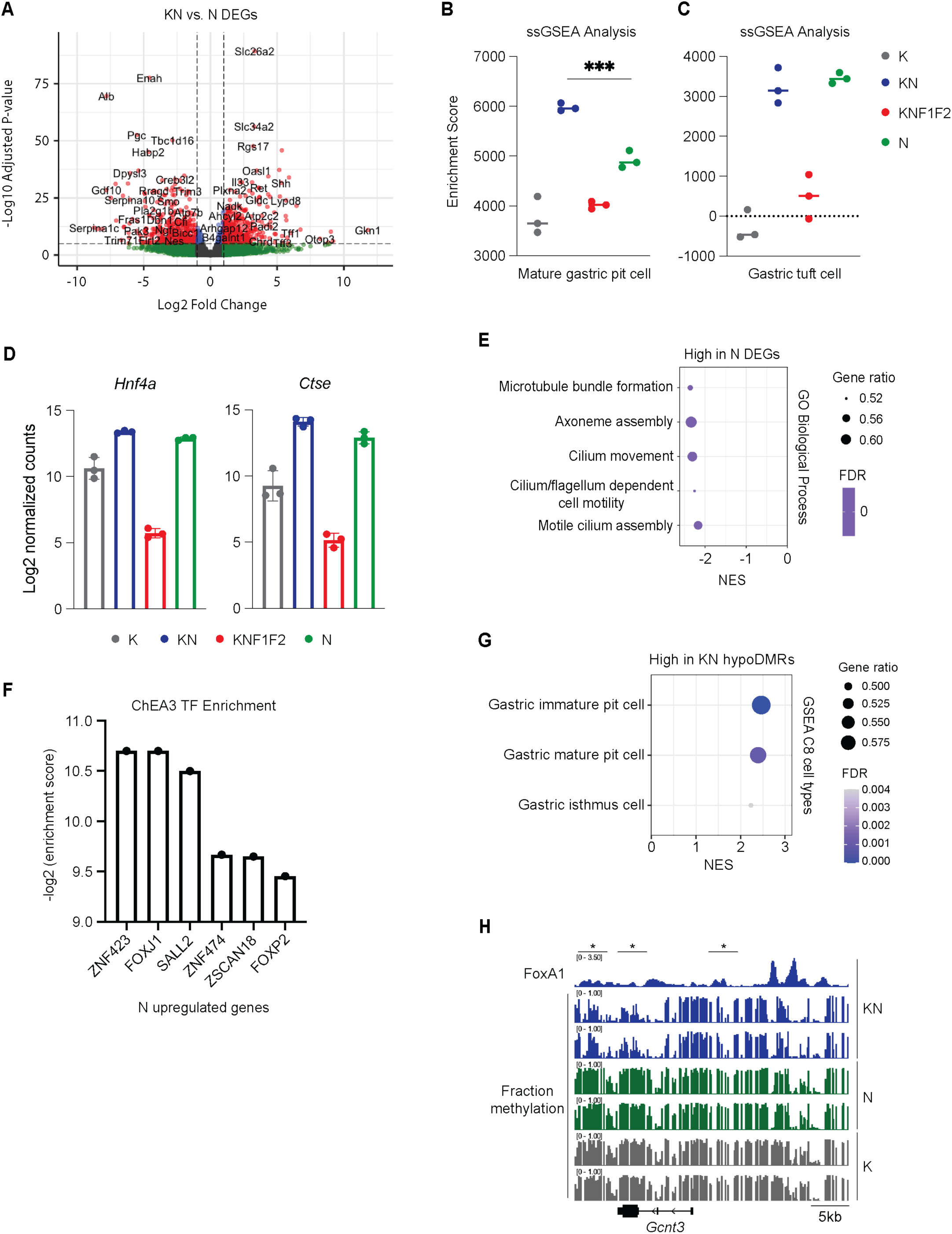
A) Volcano plot of DEGs identified between KN and N samples. Plotted are the log2 fold change values against -log_10_ adjusted *p*-value. Genes indicated by red dots have a p-value < 0.05 and log2 fold change > 1. B) Enrichment score determined by ssGSEA for mature gastric pit cell signature in K, KN, KNF1F2, and N samples. \*\*\**P*-value < 10^-3^ in KN vs. N cells, t-test. C) Enrichment score determined by ssGSEA for gastric tuft cell signature in K, KN, KNF1F2, and N samples. D) Log2 normalized counts of pan-gastric markers, *Hnf4a* and *Ctse*, in K, KN, and KNF1F2 tumors. E) GO biological processes enriched in DEGs identified between KN and N samples. F) Log2 enrichment score determined by Chea3 TF analysis for genes upregulated in N vs. KN samples. G) GSEA for cell type signatures on DMGs identified in KN-specific hypoDMRs. H) Methylation tracks of gastric pit cell gene, *Gcnt3*, in KN and N samples.

